# Slit3 Fragments Orchestrate Neurovascular Expansion and Thermogenesis in Brown Adipose Tissue

**DOI:** 10.1101/2024.09.24.613949

**Authors:** Tamires Duarte Afonso Serdan, Benjamin Frank, Heidi Cervantes, Akhil Gargey, Qiyu Tian, Daniel Hope, Chan Hee J. Choi, Anne Hoffmann, Paul Cohen, Matthias Blüher, Halil Aydin, Gary J Schwartz, Farnaz Shamsi

**Author notes:** These authors contributed equally (Benjamin Frank, Heidi Cervantes).

## Abstract

Brown adipose tissue (BAT) is an evolutionary innovation that enables placental mammals to regulate body temperature through adaptive thermogenesis. Brown adipocytes are embedded within an intricate network of blood vessels and sympathetic nerves that support their development and thermogenic function. Cold exposure activates BAT thermogenesis through the coordinated induction of brown adipogenesis, angiogenesis, and sympathetic innervation. However, how these distinct processes are coordinated remains unclear. Here, we show that fragments of Slit guidance ligand 3 (Slit3) drive crosstalk among adipocyte progenitors, endothelial cells, and sympathetic nerves. We demonstrate that adipocyte progenitors secrete Slit3, which regulates both angiogenesis and sympathetic innervation in BAT and is essential for BAT thermogenesis in vivo. Proteolytic cleavage of Slit3 generates secreted Slit3-N and Slit3-C fragments, which bind distinct receptors to stimulate angiogenesis and sympathetic innervation, respectively. We identify Plxna1 as a previously unrecognized receptor for Slit3-C and show that it is essential for sympathetic innervation and cold-induced neurite expansion in BAT. Moreover, we introduce bone morphogenetic protein 1 (Bmp1) as the first Slit protease identified in vertebrates. In summary, this work establishes a mechanistic framework for the coordinated regulation of sympathetic innervation and angiogenesis to enhance thermogenic function. The co-regulation of neurovascular expansion by distinct Slit3 fragments offers a bifurcated yet harmonized mechanism to ensure a synchronized BAT response to environmental challenges. Finally, this study provides the first evidence that adipocyte progenitors regulate tissue innervation, revealing a previously unrecognized dimension of cellular interaction within adipose tissue.

## Main

The regulation of body temperature, thermoregulation, is a fundamental homeostatic process in endothermic animals. Preserving a stable internal temperature ensures the efficiency and fidelity of all cellular reactions and is essential for survival. Brown adipose tissue (BAT) is a specialized type of adipose tissue that is primarily responsible for regulating body temperature through adaptive thermogenesis. Brown adipocytes oxidize substrates and generate heat to maintain euthermia in a cold environment^1^.

Brown adipocytes are intricately embedded in a dense network of capillaries and sympathetic nerves. The high thermogenic activity of BAT requires a high rate of blood perfusion to supply O_2_ and substrates. The sympathetic nerves also play a key role in stimulating BAT thermogenesis by releasing norepinephrine in the tissue. Norepinephrine activates adrenergic signaling in thermogenic adipocytes, resulting in enhanced thermogenesis and lipolysis^2^. Chronic cold exposure increases BAT mass by *de novo* recruitment of brown adipocytes, as well as by expanding the network of blood vessels and sympathetic nerves in the tissue. This coordinated expansion of the BAT ensures its continuous responsiveness to hormonal and neuronal stimuli and is essential for enhanced thermogenesis in cold^3,4^. However, how these distinct processes are spatiotemporally coordinated remains unclear. Furthermore, while significant progress has been made in understanding the molecular mechanisms of thermogenic activation in adipocytes, the equally crucial process of remodeling the thermogenic adipose niche remains less understood.

Cell–cell communication is vital for organismal development and function. Ligand-receptor interactions allow cells in complex tissues to coordinate their functions during development, homeostasis, and remodeling. To understand the mechanisms through which different cell types in BAT communicate and synchronize their response to cold to collectively enhance thermogenesis, we recently used single-cell transcriptomic data^5^ to construct a network of ligand-receptor interactions in BAT^6^. That study highlighted the central role of adipocyte progenitors, not merely as the source of adipocytes, but also as key communicators and versatile players in the adipose tissue microenvironment. These progenitors have been shown to contribute to a variety of processes, including extracellular matrix remodeling, immune modulation, and angiogenesis^7^.

Here, we demonstrate that Slit3 is a critical regulator of BAT neurovascular development and function. We show that Slit3 mediates crosstalk among adipocyte progenitors, endothelial cells, and sympathetic nerves, regulating both angiogenesis and innervation. Using BAT- and adipocyte progenitor-specific loss of function models, we show that the loss of Slit3 disrupts both angiogenesis and sympathetic innervation, ultimately blunting cold-induced BAT thermogenesis. We identify BMP1 as the protease responsible for Slit3 cleavage and reveal that the resulting fragments activate distinct pathways: Slit3-N promotes angiogenesis, while Slit3-C, through its newly identified receptor Plxna1, drives sympathetic innervation. This pathway demonstrates a sophisticated level of intercellular coordination whereby a single factor simultaneously drives these two distinct processes essential for thermogenic activation.

## Results

### Identification of Slit3-Robo4 signaling axis in BAT

Intercellular communication plays a critical role in coordinating tissue adaptation to external challenges. To define the role of intercellular crosstalk in cold-induced BAT thermogenesis, we previously used single-cell transcriptomic data to build a network of ligand-receptor interactions involved in the cold-induced remodeling of BAT^6^. This analysis revealed the significance of adipocyte progenitors as the major communication hub in the adipose niche^6^. This prompted us to search for ligands secreted from adipocyte progenitors that might mediate the crosstalk between adipocyte progenitors and their niche. Ligand-receptor analysis of BAT identified a new crosstalk axis in BAT involving the axon guidance molecule Slit3 and its cognate receptor, Robo4. In BAT, Slit3 is predominantly expressed in Pdgfra+ and Trpv1+ adipocyte progenitors, as well as in vascular smooth muscle cells, while its receptor, Robo4, is exclusively expressed in vascular and lymphatic endothelial cells (Figure 1a). To validate these findings, we separated the stromal vascular and adipocyte fractions of mouse BAT, sorted distinct cell populations, and measured Slit3 expression. This analysis confirmed that adipocyte progenitors are the primary source of Slit3 in BAT under both basal and cold-acclimated conditions (Figure 1b and Supplementary Figure 1a-d), whereas Robo4 is exclusively expressed in endothelial cells (Supplementary Figure 1e). To examine Slit3 protein expression in BAT and its regulation by environmental temperature, we performed Western blot analysis on BAT from mice housed at different temperatures. Slit3-FL levels were elevated in BAT of mice exposed to cold for 2 days compared to those maintained at room temperature (Figure 1c-d). These findings indicates that cold exposure enhances Slit3 protein levels in BAT.

**Figure 1.**
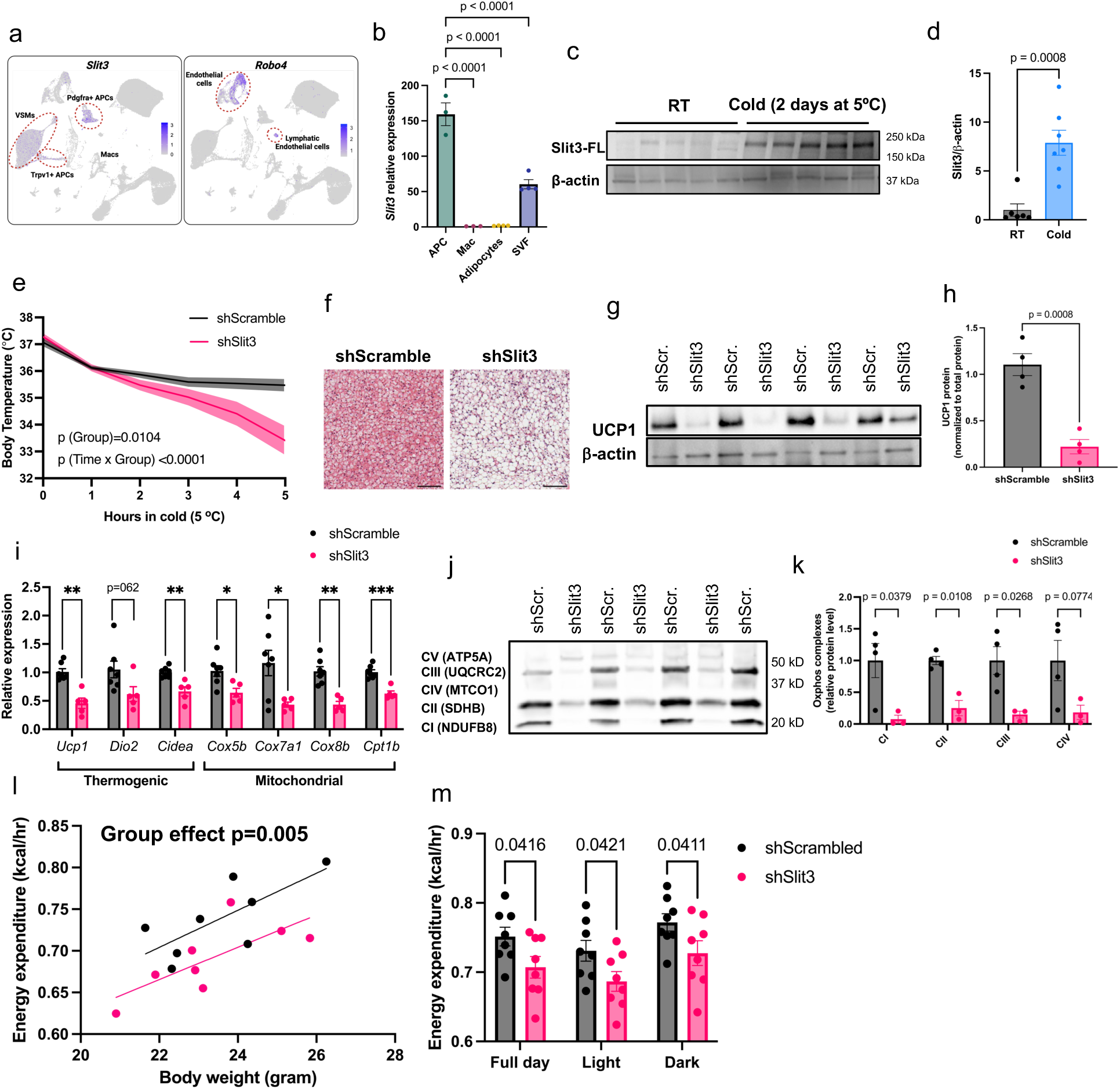
Axon Guidance Molecule Slit3 is essential for BAT thermogenesis. (a) Slit3 and Robo4 expression in scRNA-seq data from mouse BAT. (b) Slit3 expression in isolated adipocyte progenitors, macrophages, adipocytes, and the total stromal vascular fraction from mouse BAT. (c-d) Slit3 protein levels in BAT of mice housed at room temperature (RT) or cold (5°C for 2 days). (e) Cold tolerance test in AAV-shSlit3 or AAV-shScramble-injected mice. (f) Representative Hematoxylin and Eosin (H&E) staining of BAT. Scale bar = 100 µm. (g-h) UCP1 protein levels in BAT of mice after 7 days at cold (5°C). (i) Expression of thermogenic and mitochondrial genes in BAT of mice after 7 days at cold (5°C). (j-k) OxPhos complex protein levels in BAT of mice receiving AAV-shSlit3 or scramble shRNA and housed at cold (5°C) for 7 days. (l) Regression plots of energy expenditure versus total body mass in AAV-shSlit3 or AAV- shScramble-injected mice housed at 5°C. (m) Average daily energy expenditure in AAV-shSlit3 or AAV-shScramble-injected mice housed at 5°C. N = 4–7 per group (b, c-d, g-k), N = 8 per group (l, m). Data are presented as means ± SEM and analyzed by one-way ANOVA with Dunnett’s multiple comparison test (b), repeated measures ANOVA (e), unpaired two-sided Student’s t-tests (h, i, k), ANCOVA (l), and two-way ANOVA (m).

A previous study reported that Slit3 is secreted from macrophages in white adipose tissue (WAT)^8^. However, our unbiased scRNA-seq data, along with targeted expression analysis in isolated macrophages, did not detect any Slit3 expression in BAT macrophages (Figure 1a-b). Analysis of scRNA-seq datasets from mouse inguinal white adipose tissue (ingWAT) and perigonadal white adipose tissue (pgWAT), as well as human WAT^9^, revealed Slit3 expression in adipocyte progenitors, mesothelial cells, and mural cells (vascular smooth muscle cells and pericytes) (Supplementary Figure 2a-d), but no expression in macrophages in either mouse or human WAT (Supplementary Figure 2a-d). Collectively, these findings suggest that adipocyte progenitors, rather than macrophages, are the primary source of Slit3 in adipose tissue.

To assess the temporal pattern of Slit3 expression during adipogenesis, we measured SLIT3 protein levels in a mouse brown adipocyte progenitor cell line at multiple time points throughout in vitro adipogenic differentiation. SLIT3 protein levels sharply decreased by days 4 and 8 of differentiation, in contrast to the increasing UCP1 expression (Supplementary Figure 2e). These results indicate that SLIT3 expression is significantly higher in adipocyte progenitors than in mature adipocytes, where SLIT3 protein levels were minimal.

### Axon Guidance Molecule Slit3 is essential for BAT thermogenesis

To investigate Slit3 function in BAT, we used AAV-mediated shRNA delivery to knock down Slit3 expression in all Slit3-expressing cell types specifically in BAT. We confirmed a significant reduction in Slit3 transcript and protein levels in BAT, but not in WAT, of mice receiving Slit3 shRNAs (Supplementary Figure 3a-d). Mice lacking Slit3 expression in BAT exhibited severe impairments in cold-induced thermogenesis, maintaining lower core body temperature (Figure 1e) and BAT temperature (Supplementary Figure 3e) during cold exposure.

After seven days of cold exposure, analysis of BAT revealed pronounced whitening and increased lipid accumulation (Figure 1f), along with reduced expression of uncoupling protein 1 (Ucp1) (Figure 1g-h) and other thermogenic and mitochondrial genes in Slit3 knockdown mice (Figure 1i). Furthermore, Slit3 loss resulted in a marked reduction in mitochondrial electron transport chain complex levels (Figure 1j-k). Mice lacking Slit3 expression in BAT exhibited a similar reduction in Ucp1 transcript and protein levels at room temperature. However, the expression of other thermogenic and mitochondrial genes remained largely unchanged under these conditions (Supplementary Figure 4a-c).

Consistent with histological and molecular analyses, Slit3 loss in BAT significantly reduced energy expenditure, oxygen consumption, and CO₂ production in mice housed at 5°C (Figure 1l-m, Supplementary Figure 5). Importantly, these metabolic alterations occurred without changes in body weight, body composition, respiratory exchange ratio, food intake, or locomotor activity (Supplementary Figure 5). Together, these findings establish Slit3 as a critical regulator of cold-induced BAT thermogenesis and energy expenditure.

### Loss of Slit3 impairs cold-induced angiogenesis and sympathetic innervation in BAT

To investigate how the loss of Slit3 impairs cold-induced BAT thermogenesis, we first examined the cell-autonomous effects of Slit3 on adipocytes. We treated in vitro-differentiated brown adipocytes with recombinant Slit3-N and Slit3-C proteins. qPCR and immunoblot analyses showed no changes in the expression of Ucp1 or other thermogenic genes in adipocytes treated with recombinant Slit3 fragments (Supplementary Figure 6a-b). These results led us to hypothesize that Slit3 loss impairs BAT thermogenesis by disrupting the brown adipocyte niche

The expansion of BAT neurovasculature is essential for cold adaptation and enhanced thermogenesis in the cold^3,4^. Chronic cold exposure significantly increases the density of parenchymal sympathetic neurites and capillary blood vessels in BAT (Supplementary Figure 6c-f). To investigate putative mechanism(s) underlying impaired BAT thermogenesis in Slit3-deficient mice, we assessed vascularization and sympathetic innervation in BAT from Slit3 knockdown or control mice after seven days of cold exposure (5 °C). Analysis of BAT vasculature revealed that Slit3 knockdown significantly impaired angiogenesis, as indicated by a reduction in capillary density (Figure 2a-b) and a decrease in endothelial-specific transcript expression (Figure 2c).

**Figure 2.**
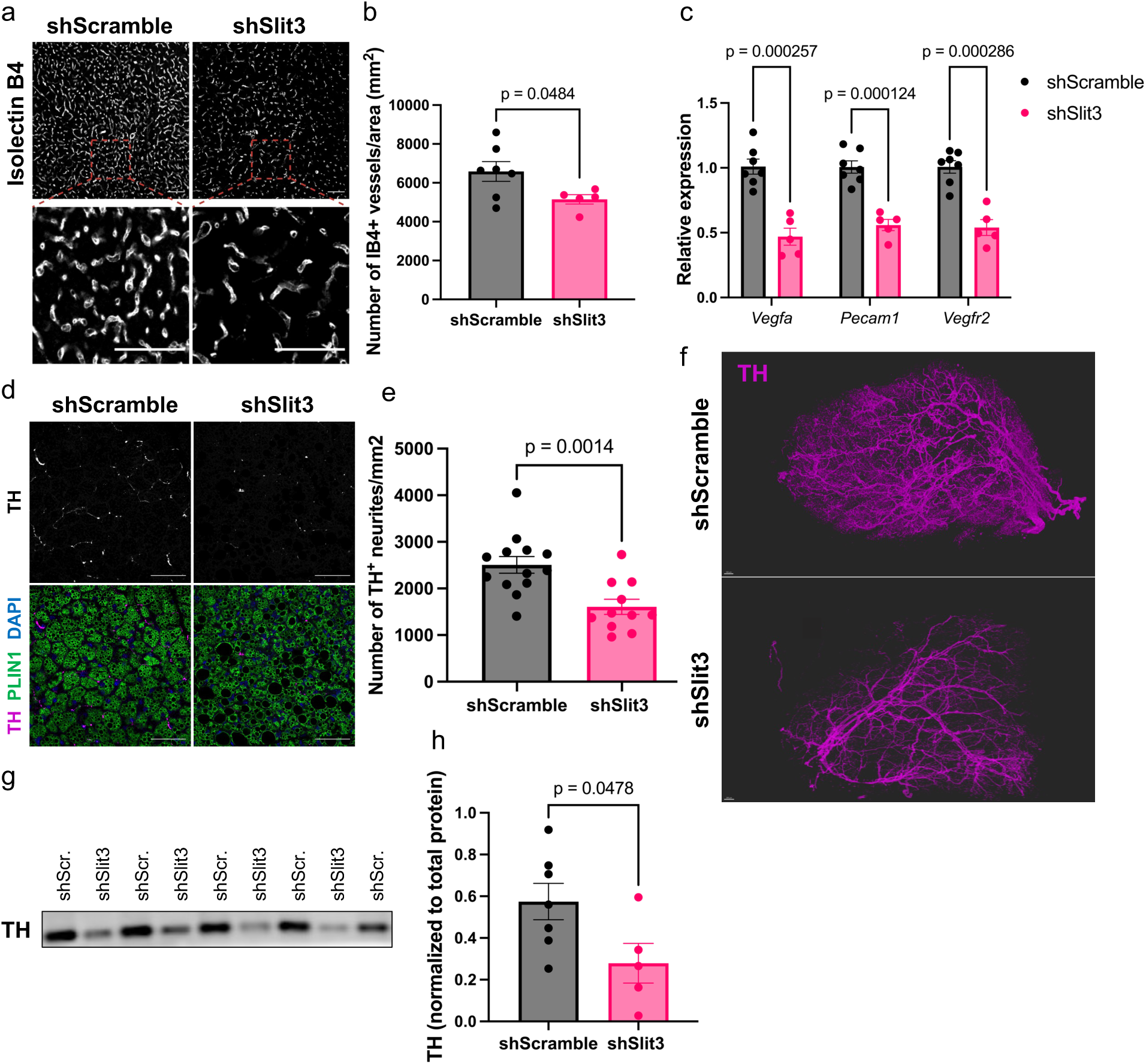
Loss of Slit3 impairs cold-induced angiogenesis and sympathetic innervation in BAT. (a) Representative images of Isolectin B4 (IB4) staining in BAT of mice receiving AAV-shSlit3 or scramble shRNA after 7 days at cold (5°C). Scale bar = 50 µm. (b) Quantification of the number of capillaries per area in BAT of mice receiving AAV-shSlit3 or scramble shRNA after 7 days at cold (5°C). (c) Expression of endothelial cell markers in BAT of mice receiving AAV-shSlit3 or scramble shRNA after 7 days at cold (5°C). (d) Representative images of TH and Plin1 staining in BAT of mice receiving AAV-shSlit3 or scramble shRNA after 7 days at cold (5°C). Scale bar = 50 µm. (e) Quantification of TH⁺ neurites per area in BAT of mice receiving AAV-shSlit3 or scramble shRNA after 7 days at cold (5°C). (f) 3-D reconstruction of TH staining in BAT of mice receiving AAV-shSlit3 or scramble shRNA after 7 days at cold (5°C). (g–h) TH protein levels in BAT of mice receiving AAV-shSlit3 or scramble shRNA after 7 days at cold (5°C). N = 5–7 per group. Data are presented as means ± SEM and analyzed by unpaired two-sided Student’s t-tests.

The loss of Slit3 also resulted in a dramatic reduction in the density of Tyrosine Hydroxylase (TH)-expressing neurites in BAT (Figure 2d-e). To assess the architecture and density of sympathetic innervation, we used the Adipo-Clear protocol^10^ to stain TH-expressing sympathetic nerves in BAT, followed by Lightsheet microscopy. 3D volumetric images of TH- positive nerves revealed a marked decline in parenchymal innervation in Slit3-deficient BAT (Figure 2f). Furthermore, total TH protein levels in BAT were significantly reduced in the absence of Slit3 (Figure 2g-h). Notably, even when animals were housed at room temperature, loss of Slit3 led to a decrease in both capillary density and sympathetic innervation in BAT (Supplementary Figure 7a-d). However, housing at cold temperatures further exacerbated these defects. These findings underscore the crucial role of Slit3 in the development and expansion of the neurovascular network in BAT, especially during adaptation to cold environment.

### Adipocyte progenitor-derived Slit3 is essential for cold-induced neurovascular expansion in BAT

To further confirm that adipocyte progenitors are the primary source of Slit3 and that progenitor-derived Slit3 is essential for cold-induced neurovascular expansion and BAT thermogenesis, we specifically deleted Slit3 in Pdgfra-expressing cells by crossing Slit3 floxed mice^11^ with an inducible Pdgfra-cre strain^12^ (Pdgfra-creERT2;Slit3^flox/flox^ or Slit3^iτιAPC^). Western blotting showed a complete loss of Slit3 protein in BAT and a marked reduction in WAT of Slit3^iτιAPC^ mice (Figure 3a-c and supplementary Figure 8a-b). Consistent with the observed phenotypes in BAT-specific Slit3 knockdown mice, both male and female Slit3^iΔAPC^ mice exhibited significant impairments in BAT thermogenesis and were cold intolerant (Figure 3d-e), highlighting the essential role of adipocyte progenitor-derived Slit3 in cold adaptation. Similar to our findings with Slit3 knockdown in BAT, Slit3^iΔAPC^ mice exposed to cold showed a significant reduction in both sympathetic neurite and capillary density in BAT (Figure f-i). Together, these results confirm that Pdgfra-expressing adipocyte progenitors are the primary source of Slit3 in BAT, and that the loss of progenitor-derived Slit3 disrupts cold-induced neurovascular expansion, leading to impaired thermogenic activation.

**Figure 3.**
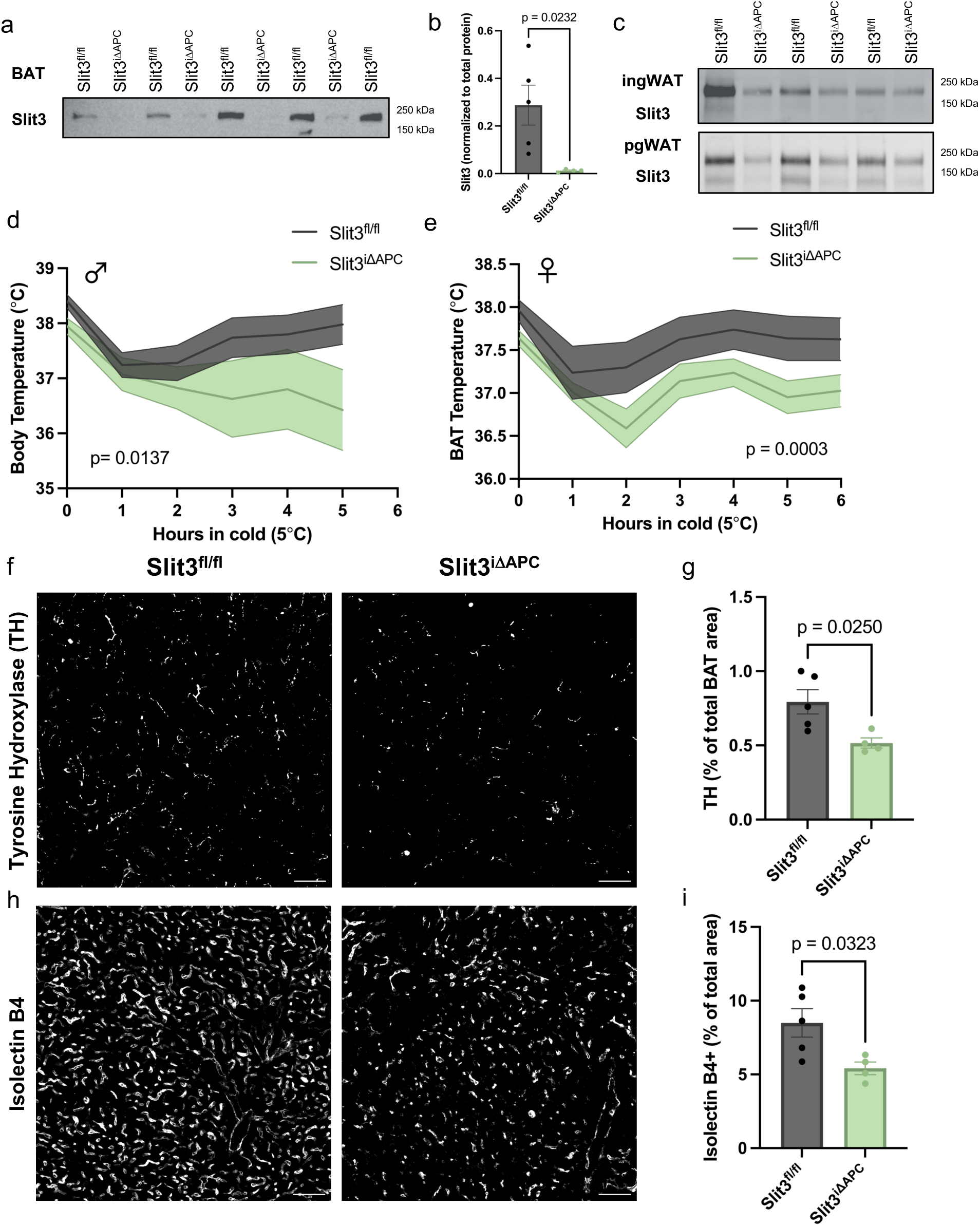
Adipocyte-specific Slit3 deletion impairs cold induced thermogenesis. (a-c) SLIT3 protein levels in BAT, ingWAT (c, top) and pgWAT (c bottom) depots of Slit3^fl/fl^ and Slit3^iΔAPC^ mice. (d-e) BAT temperature during the cold tolerance test in (g) male and (h) female Slit3^fl/fl^ and Slit3^iΔAPC^ mice. (f) Representative images of TH staining in BAT, (g) quantification of TH⁺ neurites per BAT area, (h) representative images of Isolectin B4 (IB4) staining, and (i) quantification of IB4⁺ capillaries per BAT area in BAT of Slit3^fl/fl^ and Slit3^iΔAPC^ mice after 7 days at cold (5°C). Scale bar = 50 µm. N = 4–5 per group (a–c), N = 7–9 per group (d). Data are presented as means ± SEM and analyzed by repeated measures ANOVA (d-e) and unpaired two-sided Student’s t-tests (b, g, i).

### Bmp1-mediated proteolytic cleavage of Slit3 generates two secreted ligands

Members of the Slit family are proteolytically cleaved into a large N-terminal and a shorter C-terminal fragment^13^. However, the functional implications of Slit cleavage and the distinct roles played by the full-length protein and its fragments remain unclear. Additionally, the Slit proteases in vertebrates have not been identified.

To investigate whether Slit3 is cleaved in adipocyte progenitors, we overexpressed an N- and C-terminal tagged wild-type Slit3 (SNAP-Slit3-HaloTag) as well as an uncleavable Slit3 variant (SNAP-Slit3UC-HaloTag) in a brown adipocyte progenitor cell line (Figure 4a). Using antibodies against HaloTag and SNAP-tag, we detected the full-length Slit3 protein (Slit3-FL) in the total cell lysates of cells overexpressing either the wild-type or uncleavable Slit3 constructs (Figure 4b-c). The HaloTag antibody detected two bands in the conditioned media of cells overexpressing SNAP-Slit3-HaloTag: a ∼200 kDa band corresponding to the tagged full-length Slit3 and a ∼60 kDa band matching the expected size of the tagged Slit3C-terminal fragment (Slit3-C). In contrast, the media from cells expressing the uncleavable Slit3 only contained the Slit3-FL (Figure 4b). Similarly, the SNAP-tag antibody detected both the Slit3-FL and a ∼150 kDa band corresponding to the tagged Slit3 N-terminal fragment (Slit3-N) in the media of cells overexpressing SNAP-Slit3-HaloTag (Figure 4c). A minor amount of Slit3-N, but not Slit3-C, was also detected in the total cell lysates, which contain membrane and membrane-associated proteins (Figure 4c). This finding is consistent with previous reports suggesting that the N-terminal fragment of Slits may remain associated with the plasma membrane, while the C-terminal fragment is more diffusible^13^. Collectively, these data indicate that Slit3 cleavage in adipocyte progenitors generates Slit3-N and Slit3-C fragments, and that Slit3-FL, Slit3-N, and Slit3-C fragments are secreted factors.

**Figure 4.**
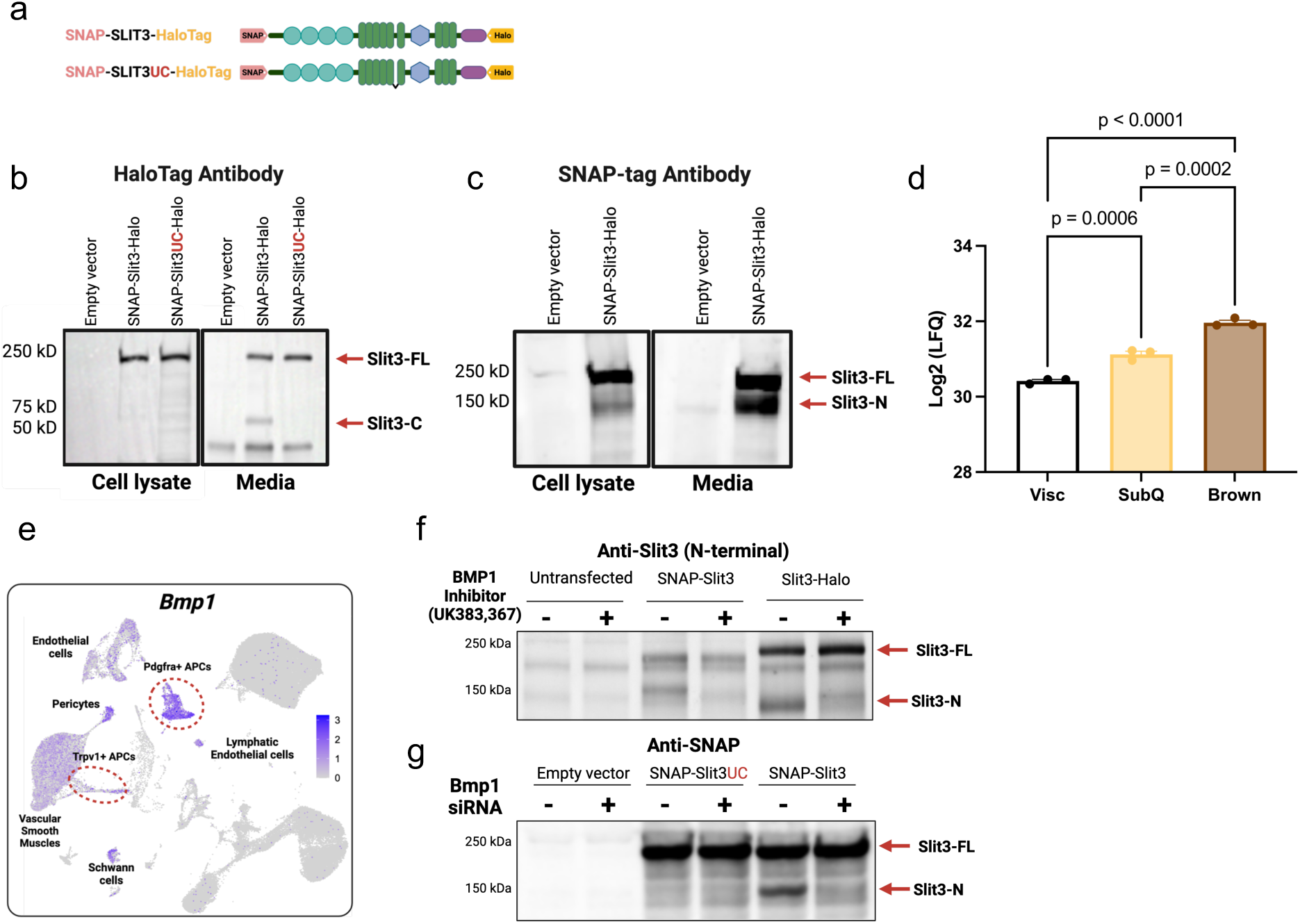
BMP1-mediated proteolytic cleavage of SLIT3 generates two secreted ligands. (a) Schematic of the tagged Slit3 transgenes. (b–c) Western blots using HaloTag (b) and SNAP-tag (c) antibodies to visualize SLIT3 fragments in cell lysates and conditioned media from adipocyte progenitors expressing the indicated plasmids. (d) Quantification of total SLIT3 peptide area in conditioned media from in vitro differentiated primary visceral (Visc), subcutaneous (SubQ), and brown adipocytes by mass spectrometry. (e) Expression of Bmp1 in adipocyte progenitors from scRNA-seq data of mouse BAT. (f) Western blot for SLIT3 in adipocyte progenitors transfected with the indicated constructs and treated with Bmp1 inhibitor UK383,367 or vehicle. (j) Western blot using SNAP-tag antibody in adipocyte progenitors transfected with siBMP1 or scrambled siRNA and overexpressing the indicated constructs. N = 3 per group. Data are presented as mean ± SEM and analyzed by one-way ANOVA with Tukey’s multiple comparisons test (d).

Moreover, analysis of a proteomics dataset of the adipocyte secretome showed the presence of Slit3 in the conditioned media of *in vitro* differentiated adipocytes derived from mouse visceral (Visc), subcutaneous inguinal white (SubQ), and interscapular BAT^14^ (Figure 4d). Slit3 was significantly more abundant in the conditioned media of brown and subcutaneous white adipocytes than visceral white adipocytes. These results indicated that Slit3 is secreted by various adipocyte types *in vitro*, with the highest abundance observed in the secretome of brown adipocytes.

A recent study suggested a role for the metalloprotease Tolkin (Tok) in Slit proteolysis in *Drosophila melanogaster*^15^. Tok is a member of the Bmp1/Tolloid family of metalloproteases. In mammals, the Bmp-1/tolloid-like family includes bone morphogenetic protein-1 (Bmp1), tolloid (Tld), and tolloid-like 1 and 2 (Tll1 and Tll2). Given the similarity in the cleavage sites among the members of the Slit family^15^, we hypothesized that the Bmp1/Tolloid proteases might be involved in Slit3 cleavage. Among the members of the Bmp1/Tolloid proteases family, only Bmp1 and Tll1 are expressed in BAT (Figure 4e and Supplementary Figure 9). Notably, Bmp1 is abundantly expressed in Pdgfra- and Trpv1-expressing adipocyte progenitors, the major cell types expressing Slit3 in BAT (Figure 4e). Bmp1 lacks growth factor activity found in other BMP family members and instead functions as a metalloprotease. To discern the role of Bmp1 in Slit3 proteolysis in adipocyte progenitors, we used a combination of pharmacological and genetic loss of function studies. We showed that the inhibition of Bmp1 activity using a small molecule inhibitor, UK383,367^16^, blocked Slit3 cleavage (Figure 4f). Similarly, knocking down Bmp1 using siRNAs prevented Slit3 cleavage (Figure 4g). Thus, we concluded that Bmp1 is responsible for the proteolytic processing of Slit3, establishing Bmp1 as the first Slit protease in vertebrates.

### Slit3 fragments promote angiogenesis and sympathetic innervation in BAT

To investigate the specific functions of Slit3 fragments in BAT, we generated AAV constructs for adipocyte-specific overexpression of full-length Slit3 (Slit3-FL), its N-terminal fragment (Slit3-N, aa 1–1116), and its C-terminal fragment (Slit3-C, aa 1117–1523) (Figure 5a). Four weeks after AAV administration in the interscapular BAT, we confirmed the specific expression of Slit3-FL, Slit3-N, and Slit3-C (Figure 5b–c). Overexpression of Slit3-FL and Slit3-C, but not Slit3-N, enhanced sympathetic innervation in BAT, as indicated by an increased number and relative area of TH-expressing sympathetic neurites (Figure 5d–f). Furthermore, overexpression of all three Slit3 variants, Slit3-FL, Slit3-N, and Slit3-C, led to increased capillary density in BAT (Figure 5g–h). Mice overexpressing Slit3-FL and Slit3-C, but not Slit3-N, also exhibited higher TH levels in BAT (Figure 5i–q and Supplementary Figure 10). Notably, Slit3-FL and Slit3-C overexpression enhanced BAT thermogenesis, as evidenced by increased Ucp1 expression and higher BAT temperatures under cold exposure (Figure 5k, 5n, 5q–r). These findings indicate that Slit3 fragments directly promote the expansion of BAT’s neurovascular network, with Slit3-C primarily driving sympathetic innervation and Slit3-N inducing angiogenesis, collectively contributing to enhanced thermogenic function.

**Figure 5.**
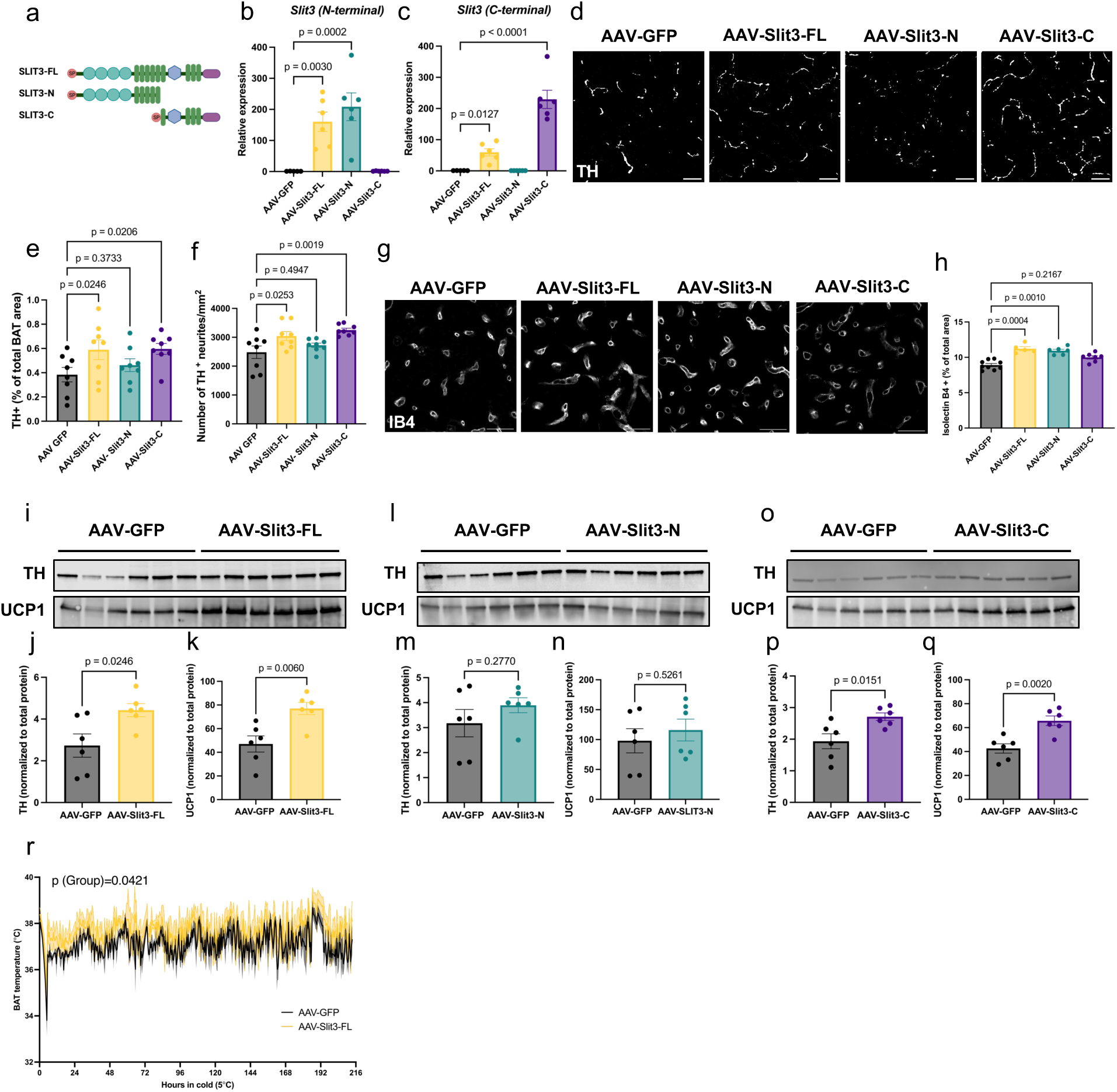
Slit3 fragments promote angiogenesis and sympathetic innervation in BAT. (a) Schematic representation of Slit3-FL, Slit3-N, and Slit3-C constructs. (b–c) Expression of Slit3 fragments in BAT measured by qPCR using primers targeting the (b) N- terminal or (c) C-terminal regions of the Slit3 transcript. (d) Representative images of TH staining in BAT from mice expressing AAV-GFP, AAV-Slit3-FL, AAV-Slit3-N, or AAV-Slit3-C after 7 days of cold exposure (5 °C). Scale bar = 25 µm. (e–f) Quantification of TH⁺ (e) area and (f) neurite density. (g) Representative images of Isolectin B4 staining in BAT from mice expressing AAV-GFP, AAV- Slit3-FL, AAV-Slit3-N, or AAV-Slit3-C after 7 days of cold exposure (5 °C). Scale bar = 50 µm. (h) Quantification of Isolectin B4⁺ capillary area in BAT. (i–q) Western blots and quantification of TH and UCP1 in BAT from mice expressing AAV-GFP, AAV-Slit3-FL (i–k), AAV-Slit3-N (l–n), or AAV-Slit3-C (o–q) after 7 days of cold exposure (5 °C). (r) BAT temperature during cold exposure in mice expressing AAV-GFP or AAV-Slit3-FL. N = 6–8 per group. Data are presented as mean ± SEM and analyzed by one-way ANOVA with Dunnett’s multiple comparisons test (b–c, e–f, h), unpaired two-sided Student’s t-test (j–k, m–n, p–q), and a mixed-effects model with Geisser-Greenhouse correction (r).

### Plxna1 is the Slit3 receptor that mediates sympathetic innervation in BAT

Full-length and N-terminal Slit fragments bind to transmembrane Robo receptors (Robo1-4)^36^, while the receptor binding and bioactivity of the C-terminal fragment (Slit-C) remains less characterized. The C-terminal of Slit2 has been shown to bind Plexin A1 (Plxna1)^17^, but the receptor(s) for Slit3-C have yet to be identified. To identify the receptors mediating Slit3 signaling in BAT, we analyzed the expression of putative Slit receptors. scRNA-seq analysis of BAT revealed that among the four Robo family members, only Robo4 is expressed in BAT, where it is exclusively localized to vascular and lymphatic endothelial cells (Figure 1a and Supplementary Figure 1e). To validate this, we co-stained BAT sections with a Robo4 antibody and Isolectin B4. We found that Robo4 was specifically localized to Isolectin B4-labeled capillaries (Figure 6a). Furthermore, double staining of BAT with Robo1 and TH antibodies revealed Robo1 expression in TH-expressing sympathetic neurites (Figure 6b). Similarly, Plxna1 and TH were co-localized in sympathetic neurites (Figure 6c).

**Figure 6.**
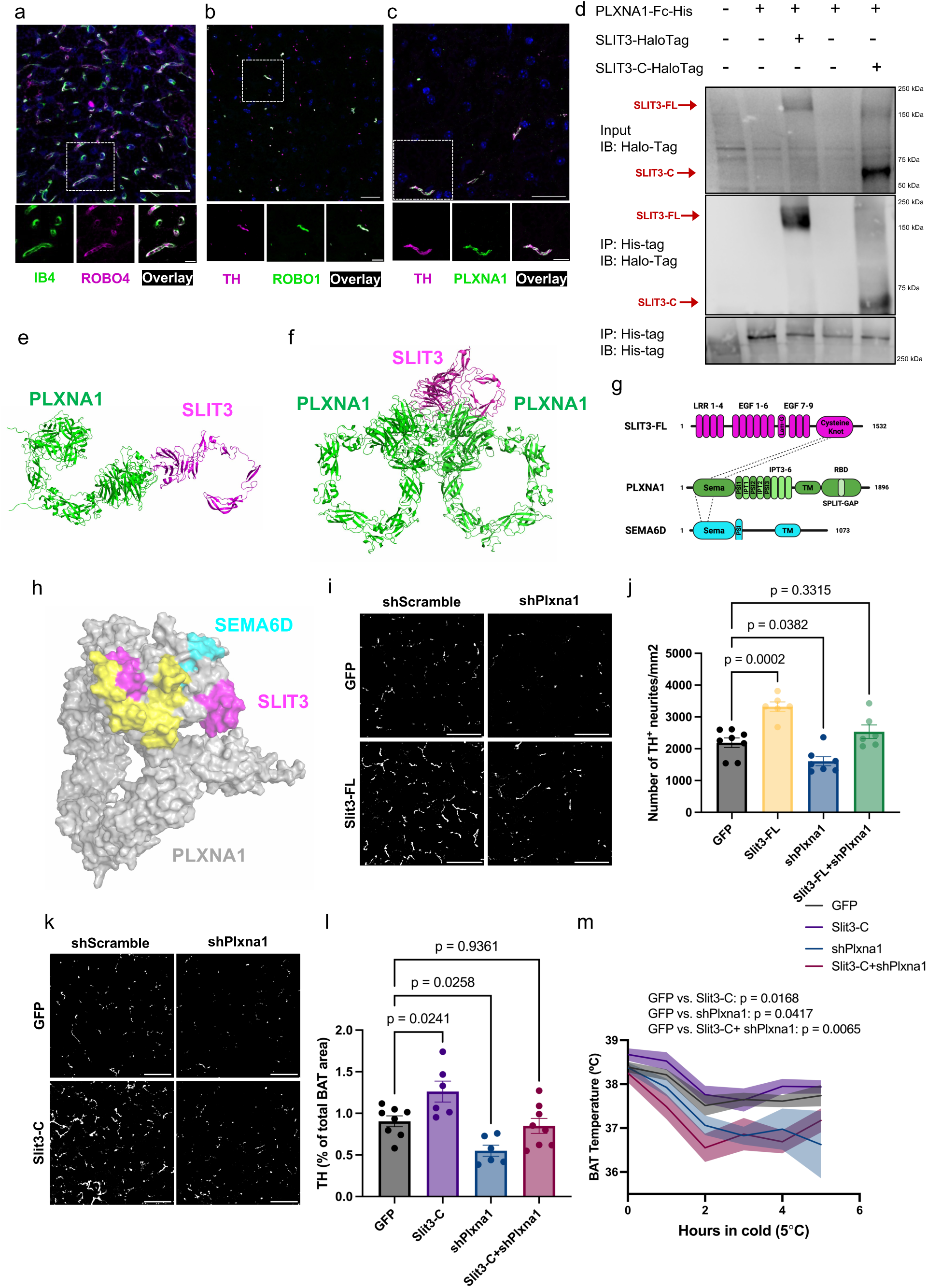
Plxna1 is the Slit3 receptor that mediates sympathetic innervation in BAT. (a–c) Immunofluorescence co-staining in BAT: (a) ROBO4 with Isolectin B4 (vascular marker), (b) ROBO1 with TH (sympathetic nerves), and (c) PLXNA1 with TH. Scale bars = 50 µm (a), 25 µm (b–c). (d) Co-immunoprecipitation of His-tagged PLXNA1 with Halo-tagged SLIT3-FL or SLIT3-C in HEK293T cells. (e-f) Ribbon representations of the top-ranked monomeric PLXNA1-SLIT3 (e) and dimeric PLXNA1 and monomeric SLIT3 (f) interaction models. (g) Domain organization of full-length human SLIT3 (magenta), PLXNA1 (green), and Semaphorin-6D (cyan). The dashed lines between SLIT3 and PLXNA1 and Semaphorin-6D (SEMA6D) and PLXNA1 indicate the binding regions between these proteins. Protein sequences used in this study are as follows: human PLXNA1 residues 27-1244 (UniProt ID: Q9UIW2), human Slit3 residues 1120-1523 (UniProt ID: O75094), human Semaphorin-6D residues 1-1073 (UniProt ID: Q8NFY4). Human SLIT3 protein comprised of an LRR: leucin-rich repeat region, an EGF-like epidermal growth factor-like domain, a Lam-G like: Agrin, Laminin, Perlecan and SLIT (ALPS) or laminin G-like module, and a C-terminal cysteine knot. SEMA6D consists of Sema and PSI domains in the extracellular region. PLXNA1 contains a Sema domain, three PSI domains, and six IPT (immunoglobulin-like, plexins, and transcription factors) domains in the extracellular domain, while a GAP (GTPase-activating protein) domain split by RBD (Rho-binding domain), termed as Split-GAP, is present in the cytoplasmic domain. (h) Human SLIT3 and SEMA6D interaction interfaces mapped on the molecular surfaces of PLXNA1 extracellular domains (residues 27-1244) AlphaFold2 model. The SLIT3 and SEMA6D binding interfaces are shown in magenta and cyan, respectively. The common binding footprint is colored in yellow. (i) Representative TH-stained images and (j) quantification of TH⁺ area in BAT of mice expressing Slit3-FL with or without shPlxna1 after 7 days of cold exposure (5 °C). Scale bar = 50 µm. (k) Representative TH-stained images and (l) quantification of TH⁺ area in BAT of mice expressing Slit3-C with or without shPlxna1 after 3 days of cold exposure (5 °C). Scale bar = 50 µm. (m) BAT temperature during the cold tolerance test in mice expressing Slit3-C with or without shPlxna1. N = 6–8 mice per group. Data are presented as mean ± SEM. Statistical analysis was performed using one-way ANOVA with Dunnett’s multiple comparisons test (f, h) and repeated measures ANOVA with Dunnett’s test (i).

Building on our finding that Slit3-C enhances sympathetic innervation (Figure 5d–f) and that Plxna1 is expressed on sympathetic neurites, we hypothesized that Plxna1 serves as the receptor for Slit3-C. To address this, we first assessed the ability of Plxna1 to interact with Slit3- FL or Slit3-C. We co-overexpressed the extracellular region of Plxna1, C-terminally fused to a 6X histidine tag, along with Slit3-FL-HaloTag or Slit3-C-HaloTag in a mouse adipocyte progenitor cell line. Using a His-tag antibody, we immunoprecipitated Plxna1 and its interacting proteins. We found that both Slit3-FL and Slit3-C co-immunoprecipitated with the extracellular region of Plxna1 (Figure 6d). These findings indicate that Plxna1 specifically interacts with the C-terminal region of Slit3 (Slit3-C), which is distinct from the N-terminal region recognized by Robo receptors^18^.

To further investigate the interaction between Plxna1 and Slit3, we used AlphaFold2 Multimer to model complexes involving the extracellular domain of Plxna1 and the C-terminal region of Slit3. Multiple structure predictions were generated based on sequence alignments and structural relaxation, with only those having average inter-protein TM (iPTM) scores >0.5 retained for further analysis. Diagnostic plots were then used to assess model quality (Supplementary Figure 11). The top-ranked models, shown in Figure 7e-f, reveal two distinct interaction modes: a monomeric Plxna1–Slit3 complex (Figure 7e) and a dimeric Plxna1 interacting with a monomeric Slit3 (Figure 7f). Figure 7g illustrates the domain organization of Plxna1, Slit3, and Sema6d—a known ligand for Plxna1—with the modeled binding regions indicated by dashed lines. The predicted binding interfaces of Slit3 and Sema6d on Plxna1’s extracellular domains (Figure 7h) suggest a direct interaction, with the Slit3 binding region in magenta, Sema6d in cyan, and a shared binding region marked in yellow. The AlphaFold2 diagnostic plots (Supplementary Figure 11) further validate the structural integrity of the models, with Predicted Aligned Error (PAE) plots for both the monomeric and dimeric complexes, along with predicted Local Distance Difference Test (pLDDT) scores, demonstrating high confidence in the predicted structures.

**Figure 7.**
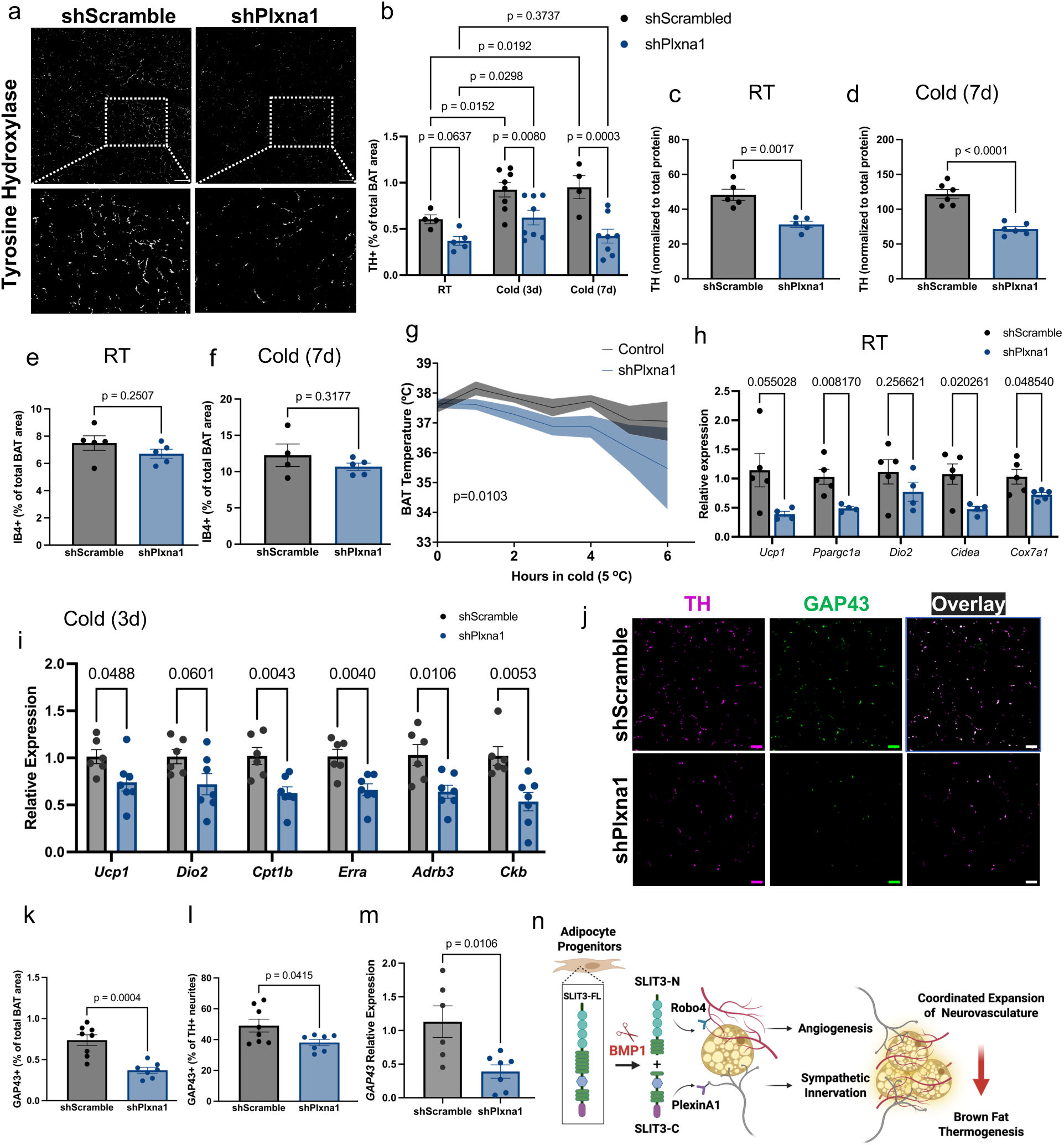
Plxna1 is essential for sympathetic innervation and cold-induced neurite expansion in BAT. All mice were injected with AAV-shPlxna1 or scramble shRNA in BAT. (a) Representative images of tyrosine hydroxylase (TH) staining in BAT. Scale bar = 50 µm. (b) Quantification of TH+ area in BAT from mice housed at room temperature (RT) or exposed to cold (5°C) for 3 or 7 days. (c–d) Total TH protein levels in BAT from mice housed at (c) RT or (d) cold (5°C) for 7 days. (e–f) Capillary density in BAT from mice housed at (e) RT or (f) cold (5°C) for 7 days. (g) BAT temperature during the cold tolerance test. (h–i) Expression of thermogenic and mitochondrial genes in BAT from mice housed at (h) RT or (i) cold (5°C) for 3 days. (j) Representative images of tyrosine hydroxylase (TH) and GAP43 staining in BAT from mice housed at cold (5°C) for 3 days. Scale bar = 20 µm. (k-l) Quantification of GAP43⁺ area (k) and percentage of TH⁺ neurites co-expressing GAP43 (l) in BAT from mice housed at 5°C for 3 days. (k) GAP43 expression in BAT from mice housed at cold (5°C) for 3 days. (n) Proposed model of BAT neurovascular remodeling regulated by Slit3 fragments. N = 4–8 mice per group. Data are presented as mean ± SEM. Statistical analysis was performed using two-way ANOVA with multiple comparisons correction (Benjamini, Krieger, and Yekutieli) for (b); unpaired, two-sided Student’s t-test for (c–f, h–i, k-m); and paired, two-sided Student’s t- test for (g).

We next examined whether Plxna1 functions as a receptor for Slit3-C in vivo and whether it is required for Slit3-mediated sympathetic innervation. To test this, we overexpressed Slit3-C in BAT with or without Plxna1 knockdown. Consistent with earlier findings (Figure 5), Slit3-FL overexpression significantly enhanced sympathetic innervation in BAT (Figure 6e–f). However, this effect was abolished when Plxna1 expression was knocked down. Similarly, Plxna1 knockdown markedly reduced the Slit3-C-induced increase in sympathetic innervation (Figure 6g–h). Notably, in the absence of Plxna1, Slit3-C overexpression failed to elevate BAT temperature in response to cold exposure (Figure 6i). Collectively, by combining biochemical assays, computational structural modeling, and in vivo studies, we provide strong and converging evidence that Plxna1 is a direct and essential receptor for the C-terminal region of Slit3, required for both sympathetic innervation and thermogenic activation in BAT.

### Plxna1 is essential for sympathetic innervation and cold-induced neurite expansion in BAT

We observed a severe reduction in sympathetic innervation of BAT and cold tolerance in mice receiving Plxna1 shRNAs alone (shPlxna1 group in Figure 6e-i), suggesting that Plxna1 is crucial for the development and cold-induced expansion of sympathetic neurites in BAT. To directly test this, we injected AAVs delivering Plxna1 shRNAs into BAT. qPCR and Western blot analysis confirmed a significant reduction of Plxna1 expression in BAT (Supplementary Figure 12a-b). Loss of Plxna1 led a pronounced decrease in sympathetic nerve density in mice housed at room temperature as well as after three and seven days of cold exposure (Figure 7a-b). This was accompanied by a reduction in total TH protein levels (Figure 7c-d and Supplementary Figure 12c-d) in BAT, while capillary density remained unchanged (Figure 7e-f). This resulted in impaired BAT thermogenesis, evidenced by reduced BAT temperature in cold conditions (Figure 7g) and a downregulation of the thermogenic gene program both at room temperature and after three days of cold acclimation (Figures 7h-i). Loss of Plxna1 reduced expression of GAP43, a key axonal growth cone protein involved in axon outgrowth and regeneration in BAT. This was reflected by a decreased GAP43⁺ neurite area and a lower proportion of sympathetic neurites expressing GAP43 (Figure 7j-l). Consistently, Gap43 mRNA levels were also reduced in BAT following Plxna1 knockdown (Figure 7m) .These findings establish Plxna1 as a key regulator of sympathetic innervation in BAT, demonstrating for the first time its essential role in both the steady-state maintenance and cold-induced expansion of sympathetic neurites in BAT (Figure 7n).

### Slit3 expression is associated with adipose tissue health and inflammation in humans

Lastly, to assess the significance of SLIT3 in human obesity, we measured its expression in human adipose tissue biopsies in abdominal subcutaneous WAT and omental visceral WAT collected from two independent cohorts of the Leipzig Obesity BioBank (LOBB). In the metabolically healthy versus unhealthy obese cohort (MHUO), we found that *SLIT3* transcript levels in human abdominal subcutaneous WAT were positively correlated with serum adiponectin concentrations (r=546, p=0.00176, N=31) and negatively regulated with the relative number of macrophages in omental visceral WAT (r=566, p<0.001, N=31) in insulin-sensitive patients (Figure 8a-b), but not in insulin-resistant patients (Figure 8c-d).

**Figure 8.**
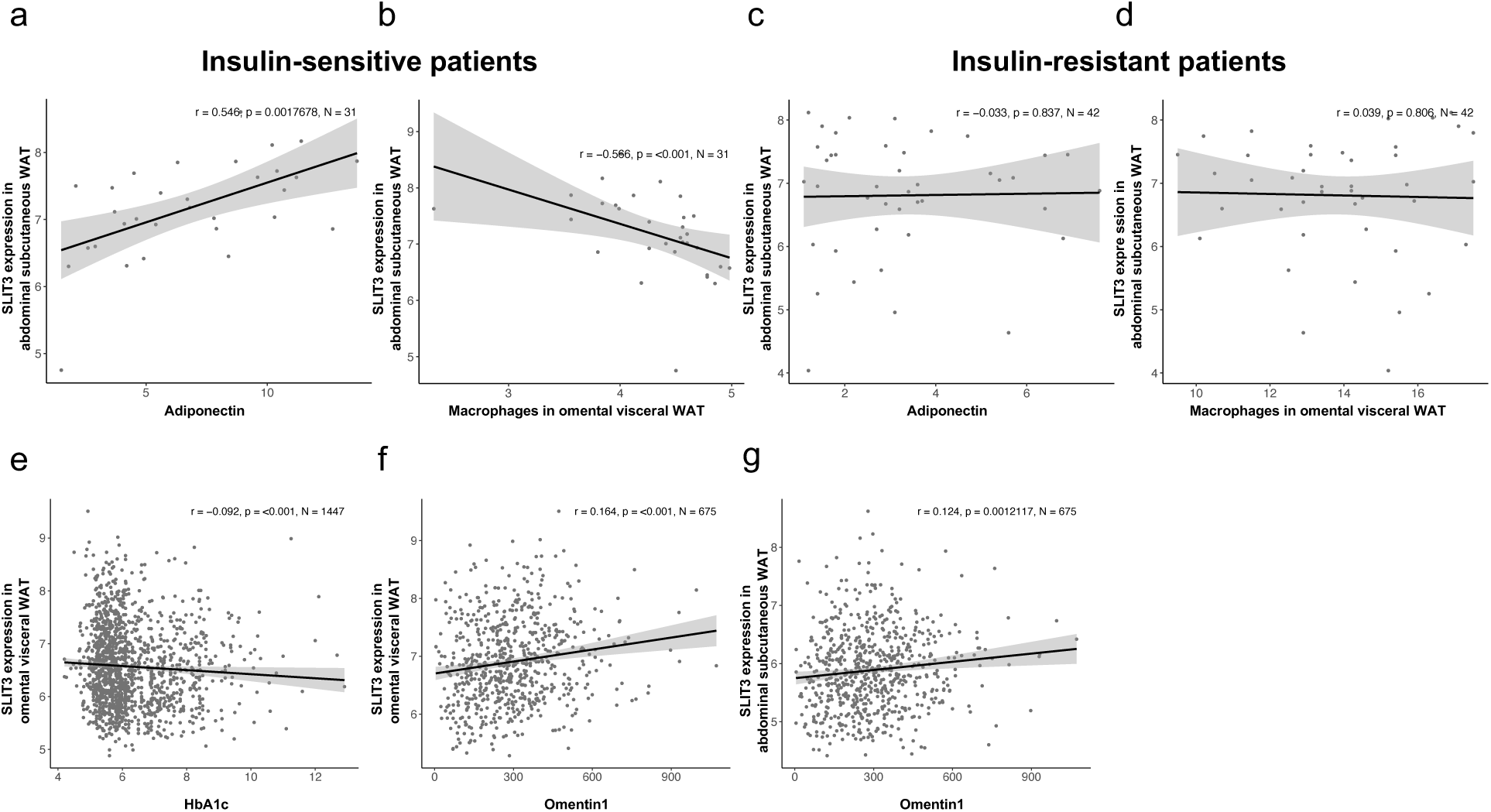
Slit3 expression is associated with adipose tissue health and inflammation in humans. Spearman correlations between SLIT3 expression in adipose tissue and metabolic or inflammatory markers in human cohorts. (a–b) In insulin-sensitive individuals, SLIT3 expression in abdominal subcutaneous WAT correlates with (a) adiponectin and (b) macrophage content in omental visceral WAT. (c–d) In insulin-resistant individuals, SLIT3 expression in abdominal subcutaneous WAT correlates with (c) adiponectin and (d) macrophages in omental visceral WAT. (e–f) In omental visceral WAT, SLIT3 expression correlates with (e) HbA1c and (f) Omentin1 levels. (g) SLIT3 expression in abdominal subcutaneous WAT correlates with Omentin1 levels. Spearman correlation coefficient (r), p-value (p), and sample size (N) are indicated on each plot. Gene expression values represent weighted trimmed mean (TMM) of log expression ratios.

Additionally, we examined *SLIT3* expression in the human cross-sectional cohort comprised of paired samples of omental visceral and abdominal subcutaneous WAT from 1,480 individuals. In this cohort, we found *SLIT3* expression in omental visceral WAT negatively correlated with Hemoglobin A1C (HbA1c) (r=0.092, p<0.001, N=1447) (Figure 8e). Additionally, *SLIT3* expression in omental visceral and abdominal subcutaneous WAT was positively associated with circulating level of the anti-inflammatory adipokine Omentin1 (r=0.164, p<0.001, N=675 and r=0.124, p=0.00121, N=675, respectively) (Figure 8f-g). Omentin1 has been shown to play important roles in glucose homeostasis, lipid metabolism, insulin resistance, and diabetes^19,20^. Collectively, these data suggest that SLIT3 may regulate adipose tissue health and inflammation in humans, potentially impacting insulin sensitivity.

## Discussion

Recent advances in technical and computational methods for studying complex tissues are providing a systematic understanding of how cellular crosstalk within the tissue microenvironment drives development, function, and disease mechanisms. Using scRNA-seq data from BAT, this study identifies Slit3 fragments as novel niche factors mediating crosstalk among adipocyte progenitors, sympathetic nerves, and blood vessels. We show that loss of Slit3 or Plxna1 in BAT significantly impairs cold-induced neurovascular expansion and adaptive thermogenesis. Our work introduces the concept that adipocyte progenitors are dynamic regulators of adipose tissue remodeling, expanding the view of their role beyond adipogenesis alone. We uncover new, molecularly specific enzymatic and receptor-mediated mechanisms that coordinate angiogenesis and sympathetic innervation—two essential processes for maintaining adipose tissue health (Figure 7n)

Despite the critical roles of Slit proteins in many processes, the biological activities of Slit fragments and the mechanisms controlling the context-dependent outcomes of Slit signaling are still largely unclear. This is in part due to the unknown identity of the Slit proteases, which prevented the separation of Slit fragment activities from that of the full-length protein. Identification of Slit proteases and the specific receptors for the Slit fragments are critical for understanding the mechanisms that control Slit signaling during tissue development, remodeling, and regeneration. In this work, we identified the metalloprotease Bmp1 as the first known vertebrate Slit protease. Through a combination of pharmacological and genetic approaches, we found that Bmp1 is responsible for Slit3 cleavage. This finding highlights Bmp1’s role in regulating the biological activities of Slit3, and potentially other Slit family members, in various contexts, including axon guidance and growth, angiogenesis, inflammatory cell chemotaxis, and tumor metastasis.

Bmp1 is a key protease that processes several extracellular matrix proteins and growth factors. It cleaves substrates such as procollagen, lysyl oxidase, and proteoglycans, which are essential for tissue integrity, including in adipose tissue^21,22^. Bmp1 also modulates the activity and availability of certain growth factors by processing their precursors, thereby influencing cell differentiation and tissue development^23^. Notably, whole-body Bmp1 knockout mice show reduced white adipose tissue mass^24^. While it’s tempting to speculate that this phenotype may be partly due to impaired Slit3 cleavage, Bmp1’s broad substrate specificity makes it difficult to draw direct conclusions from existing data.

Another key finding of our study is the identification of Plxna1 as the receptor for the C-terminal fragment of Slit3 and its essential role in regulating sympathetic innervation in BAT. Plexins are a family of single-pass transmembrane proteins that serve as receptors for Semaphorin axon guidance cues. Semaphorin-Plexin signaling regulates diverse biological processes, including axon guidance, angiogenesis, and immune responses^25^. In vertebrates, class A Plexins (PlxnAs) typically bind Class 6 Semaphorins and, together with Neuropilin co-receptors, also interact with Class 3 semaphorins^26^. Our data show that Plxna1 is required for sympathetic innervation of BAT and cold-induced induction of thermogenesis gene program. Importantly, loss of Plxna1 eliminates the pro-innervation effects of both Slit3-C and full-length Slit3. These findings challenge the conventional view that Slit proteins exclusively signal through Robo receptors and that Plexins are limited to Semaphorin signaling. Instead, we propose a previously unrecognized level of crosstalk between these two distinct ligand-receptor families. Because Plxna1 also mediates Semaphorin signaling, we cannot fully exclude the possibility that the reduced innervation observed in Plxna1-deficient BAT reflects, in part, impaired Semaphorin pathways. Future studies will be important to dissect the relative contributions of Slit3 and Semaphorin signaling through Plxna1 in regulating sympathetic innervation.

Adipose tissues are extensively innervated by a network of sympathetic and sensory nerve projections, which facilitate the transmission of information between the adipose tissue and the central nervous system. It is well-established that the sympathetic innervation of adipose tissue is critical for homeostatic control of its function and whole-body metabolism^27–29^. The plasticity of sympathetic nerves allows for altered neuronal control and adaptation to metabolic challenges. However, the mechanisms underlying the local expansion and remodeling of sympathetic neurites are not well understood. Obesity disrupts norepinephrine-mediated adipocyte lipolysis^30^, mitochondrial biogenesis^31^, and adipose tissue remodeling^32^. These impairments are attributed to dysfunctional local sympathetic innervation resulting from adipose inflammation^33^. In turn, the impaired sympathetic activity in obese adipose tissue may contribute to more inflammation and further exacerbate adipose dysfunction^34^. Therefore, preserving the density and functionality of sympathetic nerves may help safeguard adipose tissue health in obesity. To the best of our knowledge, this study is the first to demonstrate that adipocyte progenitors play a regulatory role in sympathetic innervation of adipose tissue depots, introducing a previously unexplored regulator of adipose tissue innervation.

Genome-wide association studies have found genetic variants near the SLIT3 gene to be associated with an increased risk of insulin resistance^35^ and higher BMI^36^. Additionally, a differentially methylated CpG in the SLIT3 gene in visceral adipose tissue was found to be associated with the development of type 2 diabetes in obese women^37^. Genetic variants in PLXNA1 gene are also associated with severe early onset obesity^38^. These independent findings suggest that SLIT3 and its receptor PLXNA1 may have a role in regulating metabolism in humans. Our analysis of human adipose tissue in two obesity cohorts suggest that SLIT3 signaling may contribute to improved adipose tissue health, reduced inflammation, and insulin sensitivity in human obesity. These findings highlight the need for further investigation into SLIT3-PLXNA1 signaling in the context of obesity and metabolic disease.

## Methods

### Animals

All experimental procedures involving animals were performed in compliance with all relevant ethical regulations applied to the use of small rodents and with approval by the Institutional Animal Care and Use Committees at New York University. C57BL/6J mice (stock no. 000664) were purchased from Jackson Laboratory. Pdgfra-cre;Slit3^flox/flox^ or Slit3^iτιAPC^ were generated by crossing Slit3 floxed mice^11^ with Pdgfra-creER^TM^ ^12^ (JAX strains 018280). Mice were maintained on a 12-h light-dark cycle at 22 °C and 50% relative humidity, with food and water provided ad libitum. For experiments involving cold acclimation, mice were housed at 5°C and 50% relative humidity in a controlled environmental diurnal chamber (Caron Products & Services) with free access to food and water.

### Cold tolerance test

For cold tolerance tests, mice were single-housed and placed at 5°C in a controlled environmental diurnal chamber (Caron Products & Services). Body temperature was measured with a rectal probe (Physitemp, RET3) and a reader (Physitemp, BAT-12) or using RFID transponder temperature microchips implanted under the interscapular BAT area (Unified Information Devices).

### AAV injection

6-8-week-old male mice were anesthetized with isoflurane and an incision was made above the interscapular area to expose the underlying adipose tissue. AAV particles were injected into each BAT lobe and the incision was closed with suture. Mice were allowed to recover for 3 weeks before analysis.

### Slit3 and Plxna1 shRNAs

Adeno-associated viruses (AAV8) harboring three targeting shRNA and one scramble control were purchased from VectorBuilder. The shRNA sequences are listed in Supplementary Table 1. The three shRNAs were mixed and injected into BAT at the dose of 5e+11 genomic copies per lobe.

### AAV constructs for the overexpression of Slit3 fragments in BAT

Mouse Slit3-FL, Slit3-N, and Slit3-C sequences were cloned from a cDNA clone (Origene MR225499L4) and cloned into the pAAV-hAdiponectin-W backbone^39^ using NEBuilder HiFi DNA Assembly kit using the following primers:

Slit3-FL_fwd (AACTACTCGAGgccaccATGGCCCTCGGCCGGAC),

Slit3-FL_rev

(tattcaGCGGCCGCTTAAACCTTATCGTCGTCATCCTTGTAATCgctgccGGAACACGCGCGGC A), Slit3-N_fwd (AACTACTCGAGgccaccATGGCCCTCGGCCGGAC), Slit3-N_rev (tattcaGCGGCCGCTTAAACCTTATCGTCGTCATCCTTGTAATCgctgccAACCATGGGTGGGG G), Slit3-C_fwd (tgctgccgccCTGCTACAAACCAGCCCC), Slit3-C_rev (ggttgattatcttctagagcTTACTTGTCGTCATCGTCTTTG), Signal peptide_fwd (tgattccataccagagggtcGCCACCATGGCCCTCGGC), and Signal peptide_rev (tttgtagcagGGCGGCAGCAGGGGGTCC).

pAAV-Adipoq-GFP construct^39^ was used as the control. AAV8-adipoq-Slit3FL, AAV8-adipoq- Slit3N, AAV8-adipoq-Slit3C, and AAV-Adipoq-GFP were packaged at VectorBuilder. 2e+10 AAV particles were injected into each BAT lobe.

### Fluorescence-activated Cell Sorting (FACS)

Adipocytes and the Stromal Vascular Fraction (SVF) were isolated from mouse BAT following the procedure described previously^5^. The interscapular BAT was dissected, finely minced, and digested for 45 minutes using a cocktail containing type 1 collagenase (1.5 mg ml−1; Worthington Biochemical), dispase II (2.5 U ml−1; Stemcell Technologies), and fatty acid-free BSA (2%; Gemini Bio-Products) in Hanks’ balanced salt solution (Corning Hanks’ Balanced Salt Solution, with calcium and magnesium). The resulting dissociated tissue was subsequently centrifuged at 500g and 4°C for 10 minutes. Adipocytes, located in the uppermost layer, were gently collected using a wide-mouthed transfer pipette and filtered through a 100 µm cell strainer. Brown adipocytes were allowed to float for 5 min at room temperature before they were centrifuged at 30g for 5 minutes at room temperature. This cycle was repeated three times, after which the adipocytes were immediately lysed in Trizol. For the SVF isolation, the pellet was resuspended in 10 ml of 10% FBS in DMEM, filtered through a 100 µm cell strainer into a fresh 50-ml tube, and subsequently centrifuged at 500g for 7 minutes. Red blood cells were lysed in 2 ml of sterile ammonium–chloride–potassium lysis buffer (ACK Lysing Buffer, Lonza) for 5 minutes on ice. The cells were then filtered once more through a 40-μm cell strainer, washed with 20 ml of a solution containing 10% FBS in DMEM, and centrifuged at 500g for 7 minutes. The cells were resuspended in 1 ml of Cell Staining Buffer (BioLegend) before proceeding with staining.

Cells were stained using the fluorescently conjugated antibodies as outlined in Supplementary Table 2. The cells were then incubated with the antibodies at the specified dilutions from Supplementary Table 2 for a duration of 30 minutes, followed by two rounds of washing in Cell Staining Buffer (BioLegend). Cells were sorted using an SH800 sorter using a 100 µm sorting chip (Sony Biotechnology). Debris and doublets were excluded based on forward and side scatter gating, and 7-AAD was used to exclude dead cells. Following sorting, cells were centrifuged at 300 g for 5 minutes and lysed in Trizol for subsequent RNA isolation and gene expression analysis.

### RNA isolation and quantitative reverse transcription-PCR (qRT-PCR)

RNA was isolated from cells or tissues using phenol–chloroform extraction and isopropanol precipitation. qRT PCR assays were conducted using a QuantStudio™ 5 Real-Time PCR instrument (Life Technologies Corporation) and SYBR Green (Invitrogen). Relative mRNA expression was determined using the ΔCt method and normalized to the expression of housekeeping genes Rplp0 or Tbp. Primer sequences are available in Supplementary Table 3.

### Western blotting

Cells or tissues were lysed in RIPA buffer supplemented with a protease inhibitor cocktail (cOmplete™, Sigma-Aldrich, Dallas, TX). The primary antibodies are listed in Supplementary Table 2. Primary antibodies were incubated overnight at 4 °C. HRP-coupled secondary antibodies were used at 1:3000 dilution for 1 hour at room temperature. Proteins were detected using the Amersham enhanced chemiluminescence (ECL) prime (GE healthcare, Pittsburgh, PA) using an Odyssey M Imaging System. All the original uncropped and unprocessed scans are provided in the Source Data file. For protein detection in the conditioned media, serum- free and Phenol Red-free media were collected and centrifuged at 300 g to remove cell debris. The supernatant was concentrated using Amicon™ Ultra-4 Centrifugal Filter Units (UFC801024, MilliporeSigma™).

### Immunohistochemistry

Adipose tissues were fixed in 10% formalin and embedded in paraffin. 5 µm sections were prepared and stained with the primary antibodies listed in Supplementary Table 2. Sections were deparaffinized and rehydrated, followed by an antigen retrieval step in Rodent Decloaker buffer using the Decloaking Chamber™ NxGen (BIOC Medical) in 1X rodent antigen retrieval reagent (95 °C for 45 minutes). Sections were then incubated in Sudan Black (0.3% in 70% ethanol) to reduce the autofluorescence signal. Blocking was performed in 5% BSA in PBST (0.1% Tween-20 in PBS) for 30 minutes at room temperature, followed by incubating the section in primary antibodies diluted in 1% BSA in PBST overnight at 4 °C (Supplementary Table 2). The biotinylated isolectin B4 was diluted in 5 ug/ml in a staining buffer containing 0.1mM CaCl2, 0.1mM MgCl2, and 0.1mM MnCl2. The next day, slides were washed with PBST and were incubated with appropriate fluorescently labeled secondary antibodies at a 1:200 dilution (Supplementary Table 2). Slides were stained with DAPI and mounted in a mounting medium. Images were collected on a Leica SP8 confocal microscope and processed with ImageJ.

### H&E staining

Adipose tissues were fixed in 10% formalin and embedded in paraffin. 5 µm sections were prepared and stained with hematoxylin and eosin (H&E) as previously described^40^. Slides were imaged using an Aperio slide scanner (Leica Biosystems).

### Indirect calorimetry

Mice were individually housed in metabolic cages of a TSE Phenomaster system to obtain measurements for the volume of oxygen consumption (VO2), the volume of carbon dioxide production (VCO2), respiratory exchange ratio (RER), energy expenditure, food intake, and locomotor activity. Mice had ad libitum access to drinking water and standard rodent chow. Measurements were collected first for 48 hours at room temperature, followed by the second 48 hours at 5 °C. Data were analyzed using CalR^41^.

### Dual Energy X-ray Absorptiometry

Lean and mass were quantified using a Dual Energy X-ray Absorptiometry (DEXA) scanner (Insight, Osteosys).

### Tissue clearing and imaging

Whole BAT sympathetic innervation visualization was achieved using the Adipo-Clear protocol^42^ with some modifications. One lobe of BAT from each animal was fixed in 3% glyoxal at 4°C overnight. The fixed samples were washed in PBS three times, each time for one hour. Next, the samples were subjected to a series of methanol washes: 20%, 40%, 60%, 80% methanol in H2O/0.1% Triton X-100/0.3 M glycine (B1N buffer, pH 7), and finally 100% methanol, each for 30 minutes. Subsequently, the samples underwent a triple 30-minute delipidation process with 100% dichloromethane (DCM; Sigma-Aldrich) followed by two 30- minute washes in 100% methanol. After delipidation, the tissues were bleached overnight at 4°C with 5% H2O2 in methanol (1 volume of 30% H2O2 to 5 volumes of methanol). The rehydration process followed a reversed methanol/B1N buffer series: 80%, 60%, 40%, 20% methanol in B1N buffer, each step lasting 30 minutes. All the above steps were conducted at 4°C with continuous shaking. Subsequently, samples were washed in B1N buffer twice, each time for 30 minutes, followed by two 1-hour wash in PBS/0.1% Triton X-100/0.05% Tween 20/2 µg/ml heparin (PTwH buffer), before initiating the staining procedure. The samples were incubated with anti-tyrosine hydroxylase antibody (1:200, AB1542, Millipore Sigma) in PTxwH for 4 days. After the primary antibody incubation, the samples underwent a series of PTxwH washes lasting 5 min, 10 min, 15 min, 30 min, 1 hr, 2 hr, 4 hr, and overnight, and then were incubated in donkey anti-sheep IgG Alexa Fluor 647 (1:200, A-21099, Thermo Fisher) in PTxwH for 4 days. Finally, the samples were washed in PTwH for 5 min, 10 min, 15 min, 30 min, 1 hr, 2 hr, 4 hr, and overnight. Finally, the samples were dehydrated using a methanol/H2O series (25%, 50%, 75%, 100%, 100%) for 30 minutes at each step at room temperature. Following dehydration, samples were incubated with 100% DCM for 30 minutes twice, followed by an overnight clearing step in dibenzyl ether (DBE; Sigma-Aldrich). The samples were then stored at room temperature in the dark until imaging. Representative samples were imaged by a blinded experimenter using a Zeiss Lightsheet Z.1 Microscope with two 1920 × 1920-pixel sCMOS cameras. Three-dimensional reconstructions were generated using Imaris (Bitplane).

### Cloning of SNAP- and Halo-tagged Slit3 Plasmids

pRP[Exp]-CMV>[SNAP-Slit3-HaloTag], pRP[Exp]-CMV>[Slit3-FL]-HaloTag, and pRP[Exp]-CMV>[Slit3-C-HaloTag] constructs were generated by VectorBuilder. The N-terminal tag was added immediately downstream of the signal peptide (aa 2-33,

GCCCTCGGCCGGACCGGGGCCGGCGCCGCTGTGCGCGCCCGCCTGGCGCTGGGCTTGGCGCTTGCGAGCATCCTGAGCGGACCCCCTGCTGCCGCC).

The ORF in the pRP[Exp]-CMV>[SNAP-Slit3UC-Halo] was generated by the deletion of a 27 bp sequence encoding the Slit cleavage site (CCCACCCATGGTTCTGCTACAAACCAG, PPPMVLLQ) from Slit3-FL cDNA.

### Transient transfection

Immortalized brown preadipocytes were transfected with Slit3 overexpression plasmids, Bmp1 or scramble siRNA (Dharmacon GE) using Lipofectamine™ 3000 or Lipofectamine™ RNAiMAX Transfection Reagents (Invitrogen™) following the manufacturer’s instructions.

### Bmp1 inhibitor treatment

Cells were treated with the Bmp1 inhibitor, UK-383,367 (PZ0156- 5MG, Sigma Aldrich) at the final concentration of 2.5 μM.

### AlphaFold Multimer Analysis

AlphaFold2 Multimer was used to predict the protein-protein interactions between the extracellular domains of human PLXNA1 and the C-terminal region of the Slit3 protein as described previously^43,44^. Briefly, a local graphics processing unit (GPU) cluster using MMseqs (git@92deb92) was utilized for local Multiple Sequence Alignment (MSA) creation^45^, and ColabFold (git@7227d4c) was executed for structure prediction with 5 models per prediction, including structure relaxation^46^. Predictions with an average iPTM score of >0.5 were ranked, and diagnostic plots (PAE plot, pLDDT plot, and sequence coverage) were manually inspected. The generated three-dimensional (3D) models of protein-protein complexes were analyzed using UCSF Chimera^47^ and the PyMOL Molecular Graphics System, Version 3.0 Schrödinger, LLC.

### Human data

The human data used in this research were sourced from the Leipzig Obesity Biobank (LOBB, https://www.helmholtz-munich.de/en/hi-mag/cohort/leipzig-obesity-bio-bank-lobb), which comprises paired samples of abdominal subcutaneous and omental visceral adipose tissue. The metabolically healthy versus unhealthy obese cohort (MHO/MUO) comprises paired samples of omental visceral and abdominal subcutaneous tissues from 31 insulin-sensitive patients (71% female; age: 38.8 ± 11.1 years old; BMI: 45.9 ± 6.9 kg/m²; fasting plasma glucose: 5.2 ± 0.2 mmol/l; fasting plasma insulin: 27.9 ± 13.5 pmol/l) and 42 insulin-resistant patients (71.43% female; age: 47.2 ± 7.7 years old; BMI: 47.3 ± 8.1 kg/m²; fasting plasma glucose: 5.7 ± 0.3 mmol/l; fasting plasma insulin: 113.7 ± 45.7 pmol/l). The cross-sectional cohort (CSC) comprises 1,479 individuals, categorized as either normal/overweight (N = 31; 52% women; age: 55.8 ± 13.4 years; BMI: 25.7 ± 2.7 kg/m²) or obese (N = 1,448; 71% women; age: 46.9 ± 11.7 years; BMI: 49.2 ± 8.3 kg/m²). Adipose tissue samples were collected during elective laparoscopic abdominal surgeries, following established protocols^48,49^. Body composition and metabolic parameters were assessed using standardized techniques as described previously^50,51^. The study was approved by the Ethics Committee of the University of Leipzig (approval number: 159-12- 21052012) and adhered to the principles outlined in the Declaration of Helsinki. All participants provided written informed consent before being included in the study. Exclusion criteria included individuals under 18 years of age, those with chronic substance or alcohol abuse, smoking within the 12 months prior to surgery, acute inflammatory conditions, concurrent use of glitazones, end-stage malignancies, weight loss greater than 3% in the three months leading up to surgery, uncontrolled thyroid disorders, and Cushing’s disease.

Ribosomal RNA-depleted RNA sequencing data we generated following the SMARTseq protocol^52^. The libraries were sequenced as single-end reads on a Novaseq 6000 (Illumina, San Diego, CA, USA) at the Functional Genomics Center Zurich, Switzerland. The preprocessing procedures were conducted as previously described^53^. In brief, adapter and quality-trimmed reads were aligned to the human reference genome (assembly GRCh38.p13, GENCODE release 32), and gene-level expression quantification was performed using Kallisto^54^ v0.48. For samples with read counts exceeding 20 million, we downsampled them to 20 million reads utilizing the R package ezRun v3.14.1 (https://github.com/uzh/ezRun, accessed on April 27, 2023). The data normalization was carried out using a weighted trimmed mean (TMM) of the log expression ratios, with adjustments made for age, sex, and transcript integrity numbers (TINs). Analyses were conducted in R v4.3.1 (www.R-project.org). Spearman coefficient was used to assess the correlation between *SLIT3* expression and metabolic parameters.

### Data reporting

No statistical methods were used to predetermine the sample size. Experiments were not randomized. All imaging and quantifications were performed in a blinded manner. Statistical analyses were performed using GraphPad Prism.

### Data availability

The data supporting the findings of this study are available within the paper and its supplementary information files. The human RNA-seq data from the LOBB have not been deposited in a public repository due to restrictions imposed by patient consent but can be obtained from Matthias Blüher upon request.

## Acknowledgment

This work was supported in part by the US National Institutes of Health (grants K01DK125608, R03DK135786, R01DK136724), an award from The G. Harold and Leila Y. Mathers Charitable Foundation, the American Heart Association Career Development Award, and a grant from Einstein-Mount Sinai Diabetes Center (to F.S.) and by National Institutes of Health grant RC2DK129961 (to P.C.). The Albert Einstein Animal Physiology Core is supported by the NIH grant DK20541. M.B. received funding from grants from the DFG project number 209933838 – SFB 1052 (project B1) and by Deutsches Zentrum für Diabetesforschung (DZD, Grant: 82DZD00601). The Slit3 floxed strain was kindly provided by Dr. Matthew Greenblatt. We thank Adam Mar and Begona Gamallo-Lana at the NYU Langone Rodent Behavior Laboratory (RRID: SCR_017942) for providing technical assistance, Experimental Pathology Research Laboratory (RRID:SCR_017928), Preclinical Imaging Laboratory (RRID:SCR_017937), and Microscopy Laboratory (RRID: SCR_017934). The microscopy shared resource is partially supported by Cancer Center Support Grant P30CA016087. We thank Christian Wolfrum’s group for contributing to the LOBB human adipose tissue RNA sequencing.

## Author Contributions

T.D.S. and F.S. conceptualized the work and designed research. T.D.S., B.F., H.C., Q.T., and D.H. performed research and analyzed data. A.G. and H.A. performed the AlphaFold Multimer Analysis. C.H.J.C. and P.C. provided the adipocyte secretome data. A.H. performed RNA sequencing and preprocessing of the LOBB data. M.B. provided the data from the LOBB cohorts. G.J.S helped in designing the experiments and provided feedback. F.S. wrote the manuscript with input from all authors.

---

**Supplementary Figure 1. Related to Figure 1.**
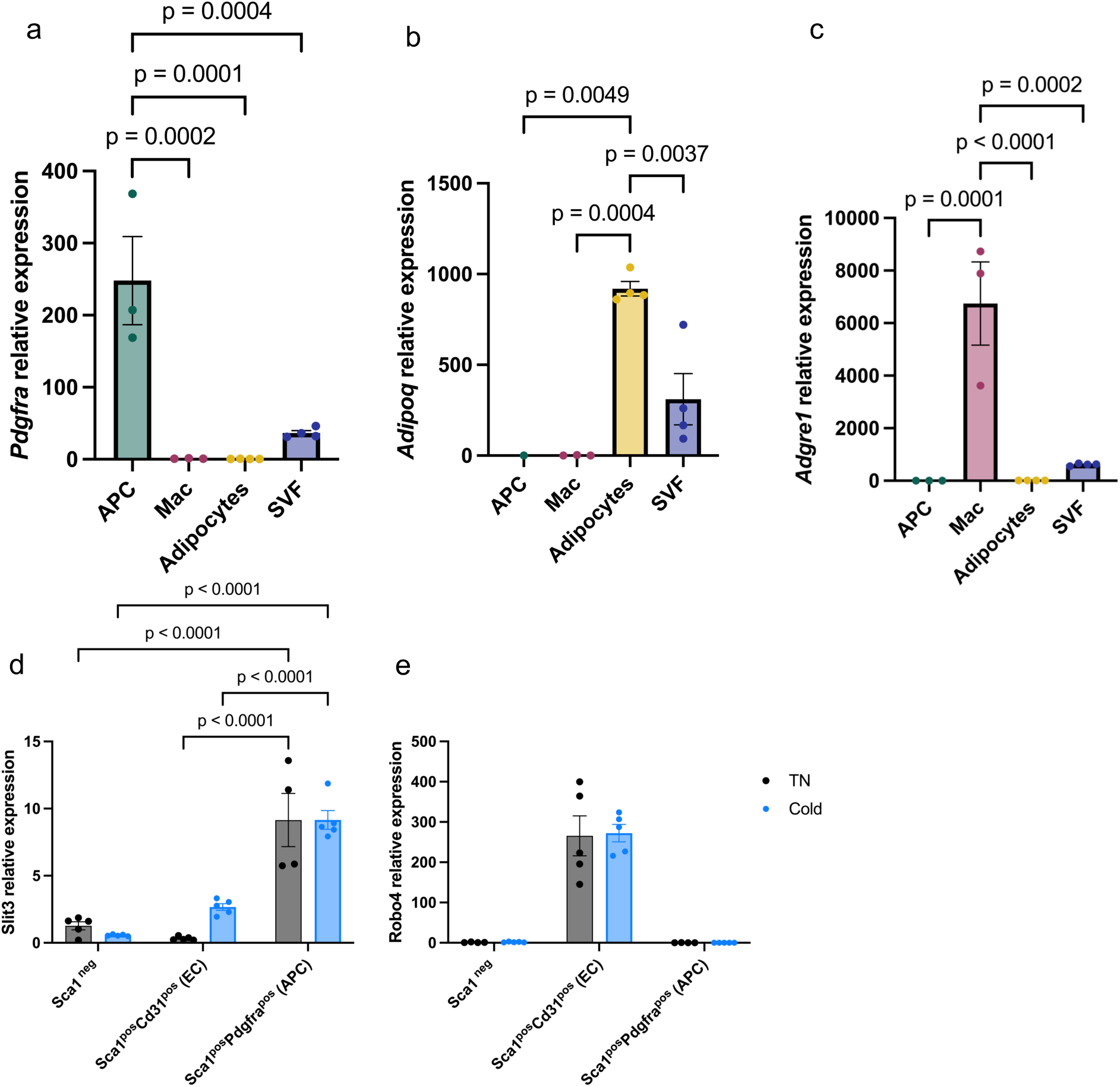
(a–c) Expression of *Pdgfra*, *Adipoq*, and *Adgre1* in isolated adipocyte progenitors, macrophages, adipocytes, and total stromal vascular fraction (SVF) from mouse BAT. (d–e) Expression of *Slit3* and *Robo4* in Sca1⁻, Sca1⁺Cd31⁺, and Sca1⁺Pdgfra⁺ cells isolated from BAT of mice housed at thermoneutrality (30 °C) or cold (5 °C) for 7 days. N = 3–5 per group. Data are presented as mean ± SEM and analyzed by one-way ANOVA with Dunnett’s multiple comparisons test (a–c) and two-way ANOVA with Tukey’s multiple comparisons test (d–e).

**Supplementary Figure 2. Related to Figure 1.**
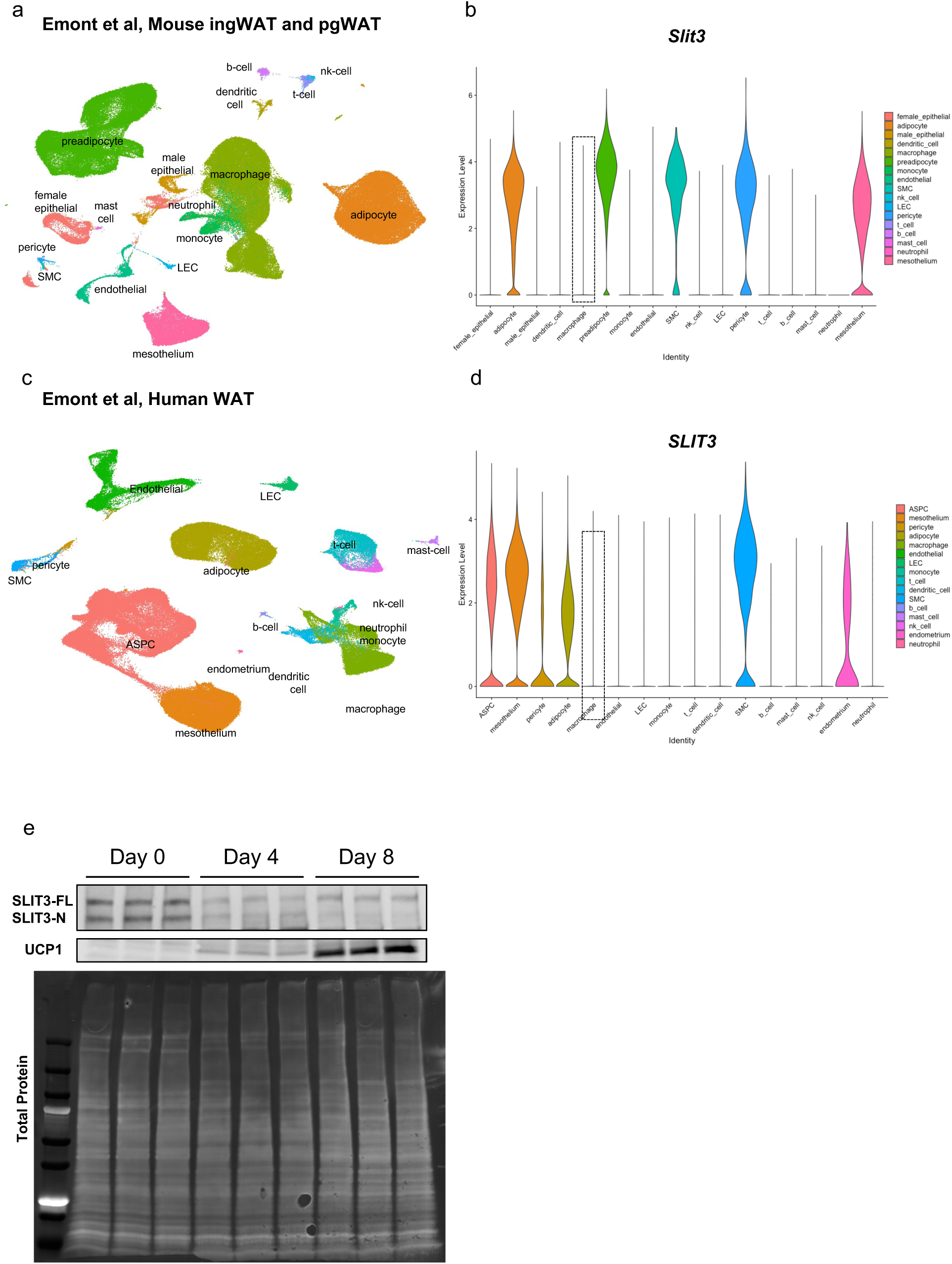
(a–b) UMAP of unsupervised clustering of nuclei from mouse ingWAT and pgWAT (a) and violin plot showing Slit3 transcript expression across clusters (b). (c–d) UMAP of unsupervised clustering of nuclei from human WAT (c) and violin plot showing SLIT3 transcript expression across clusters (d). (e) SLIT3, UCP1, and total protein levels in SVF-derived brown adipocytes at day 0, day 4, and day 8 of differentiation.

**Supplementary Figure 3. Related to Figure 1.**
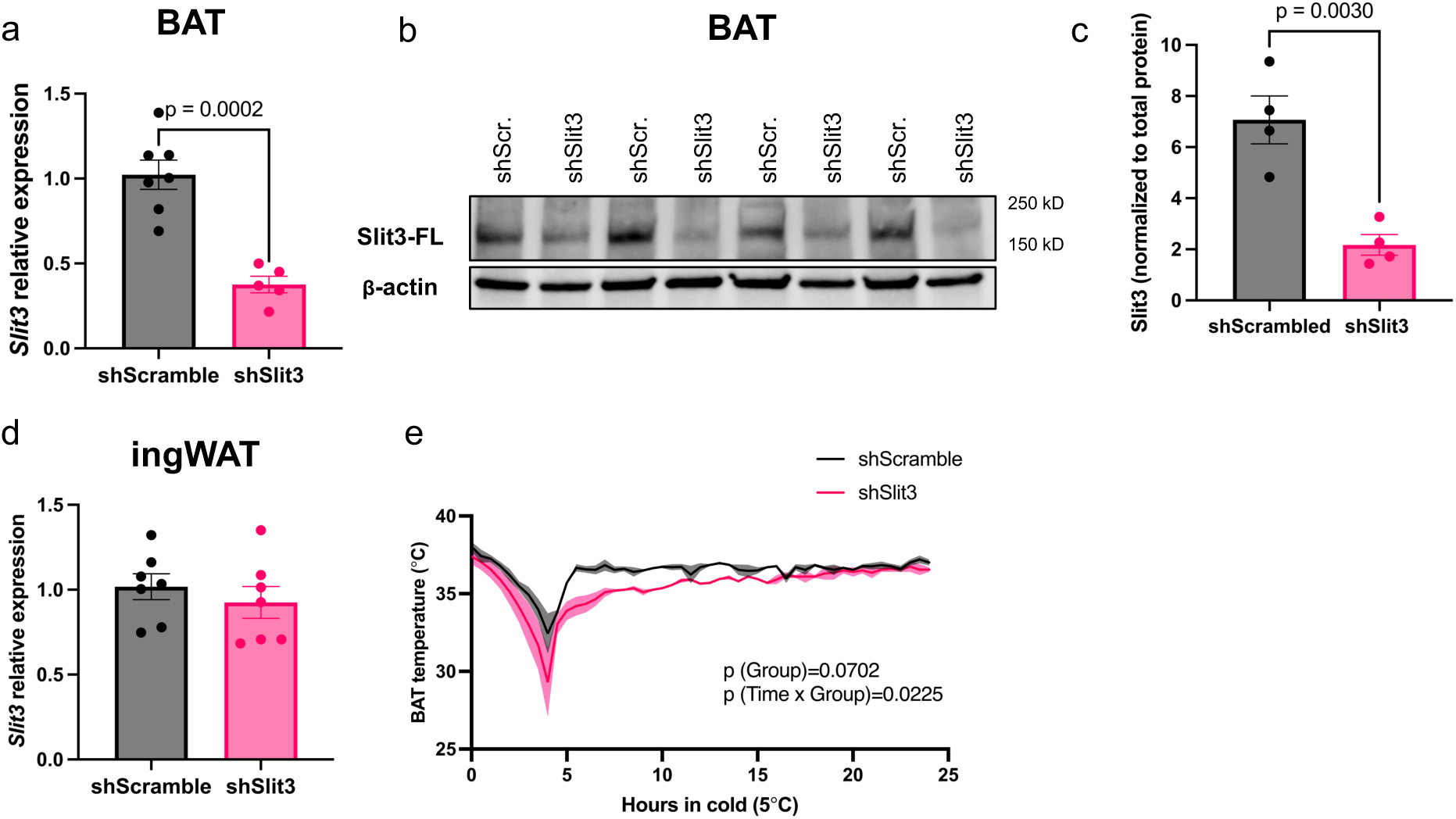
(a) Slit3 transcript levels and (b–c) protein levels in BAT, (d) Slit3 transcript levels in inguinal WAT (ingWAT), and (e) BAT temperature in mice injected with AAV-shSlit3 or scramble shRNA and housed at cold (5 °C) for 7 days. N = 5–7 mice per group. Data are presented as mean ± SEM and analyzed by unpaired two-sided Student’s t-tests (a, c) and repeated measures ANOVA (d).

**Supplementary Figure 4. Related to Figure 1.**
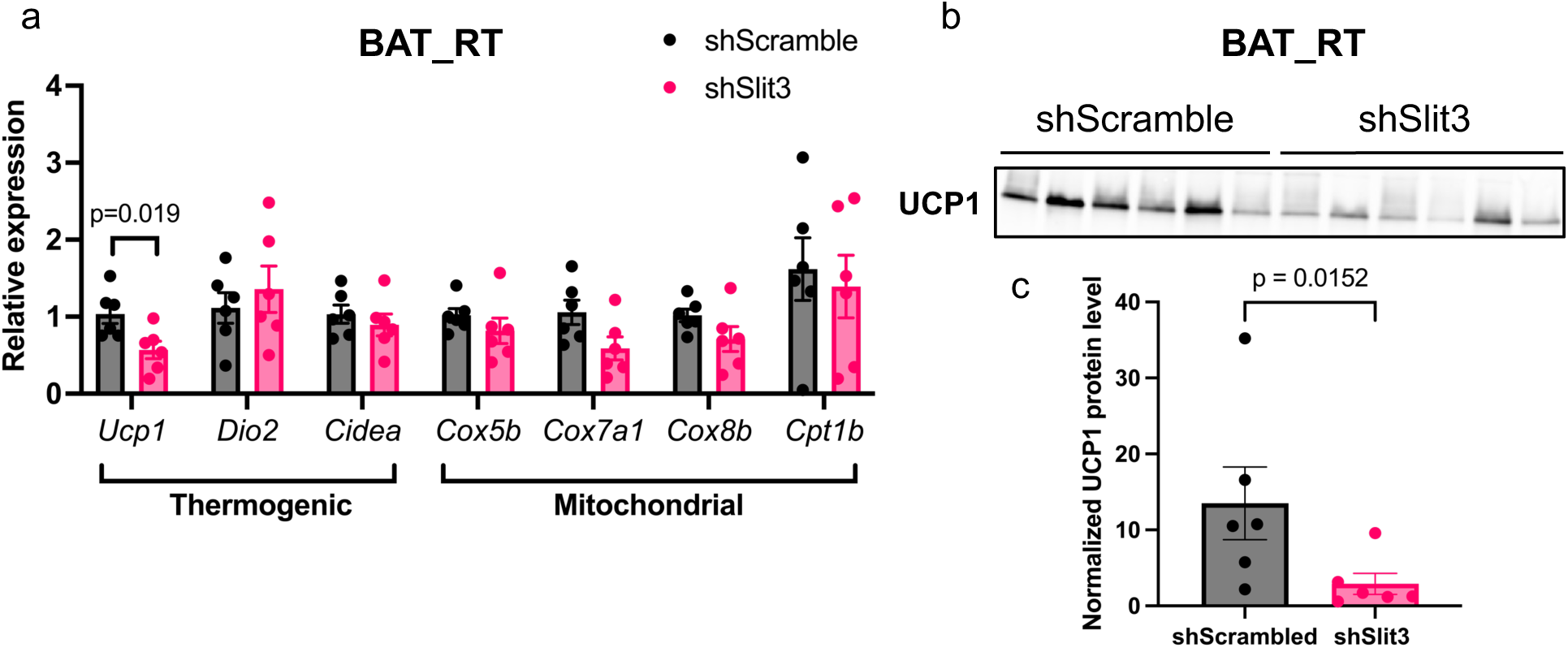
(a) Expression of thermogenic and mitochondrial genes, and (b) UCP1 protein levels in BAT of mice injected with AAV-shSlit3 or scramble shRNA and housed at room temperature. N = 6 per group. Data are presented as mean ± SEM and analyzed by unpaired two-sided Student’s t-test (a and c).

**Supplementary Figure 5. Related to Figure 1.**
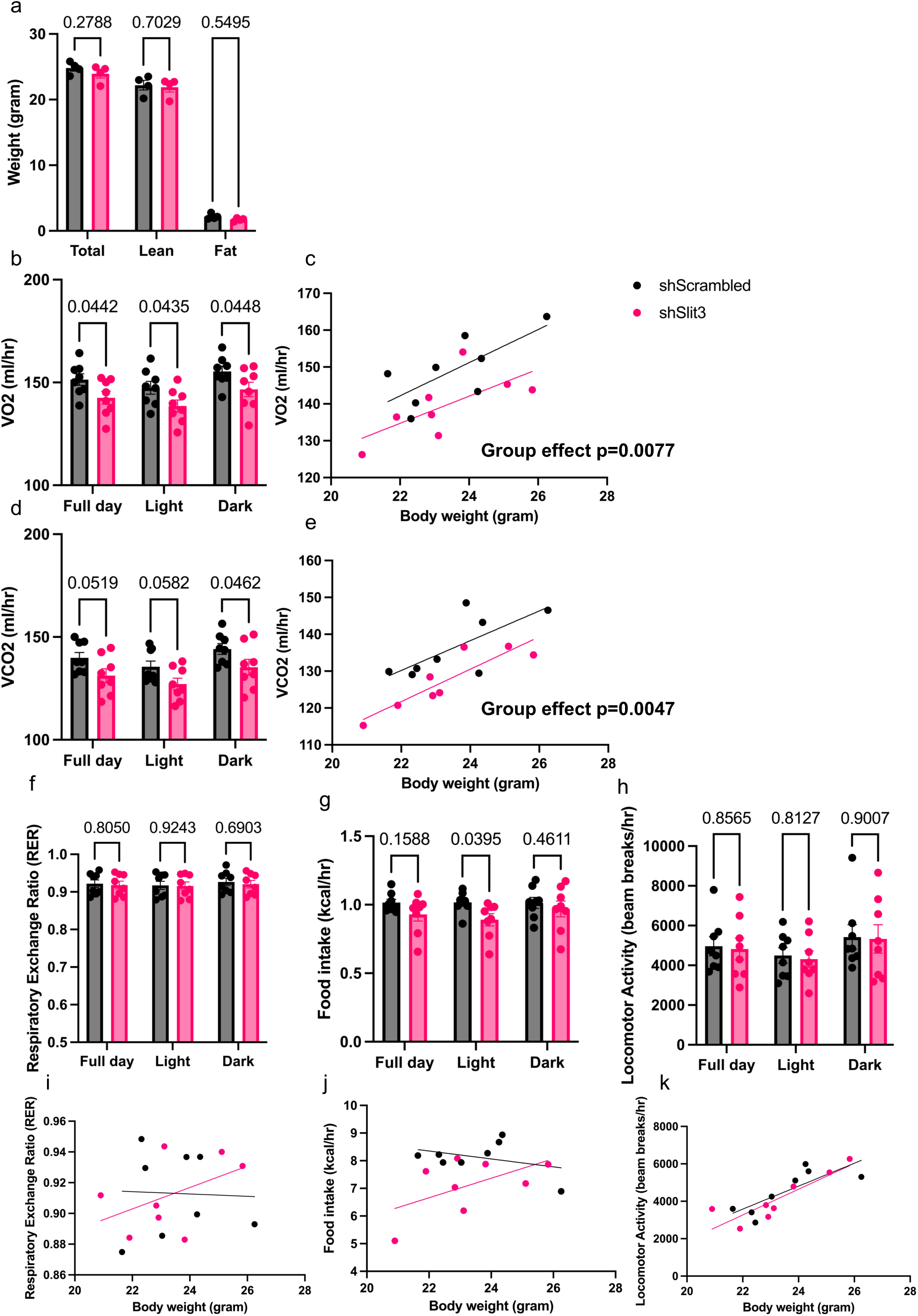
(a) Body composition analysis in mice injected with AAV-shSlit3 or scramble shRNA and housed at cold (5 °C) for 3 days. (b, d, f–h) Average hourly rates of VO₂, VCO₂, respiratory exchange ratio (RER), food intake, and locomotor activity. (c, e, i–k) Regression plots of VO₂ (c), VCO₂ (e), RER (i), food intake (j), and locomotor activity (k) versus total body mass. N = 8 mice per group. Data are presented as mean ± SEM and analyzed by unpaired two-sided Student’s t-test (a), two-way ANOVA (b, d, f–h), and ANCOVA (c, e).

**Supplementary Figure 6. Related to Figure 2.**
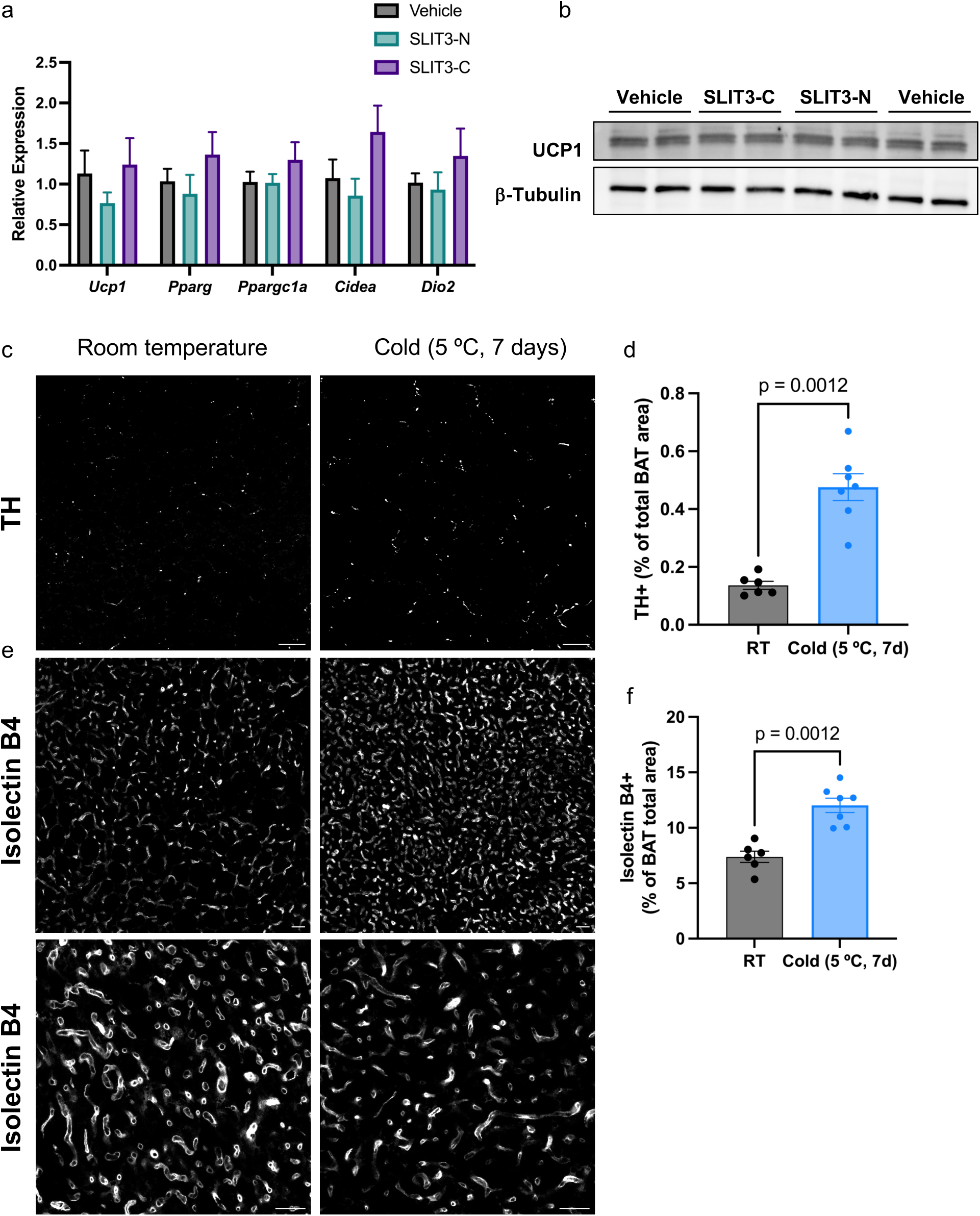
(a) Expression of thermogenic genes in in vitro differentiated brown adipocytes treated with recombinant Slit3-C (100 ng/ml), Slit3-N (100 ng/ml), or vehicle for 48 hours. (b) UCP1 protein levels in in vitro differentiated brown adipocytes treated with recombinant Slit3- C, Slit3-N, or vehicle for 48 hours. (c) Representative images of TH and Plin1 staining and (d) quantification of TH+ neurites per area in BAT of mice housed at room temperature and cold (5°C) for 7 days. Scale bar = 50 µm. (e) Representative images of Isolectin B4 staining and (f) quantification of the number of capillaries per area in BAT housed at room temperature and cold (5°C) for 7 days. Scale bar = 50 µm. N = 5–7 per group. Data are presented as means ± SEM and analyzed by unpaired two-sided Student’s t-tests (d and f).

**Supplementary Figure 7. Related to Figure 2.**
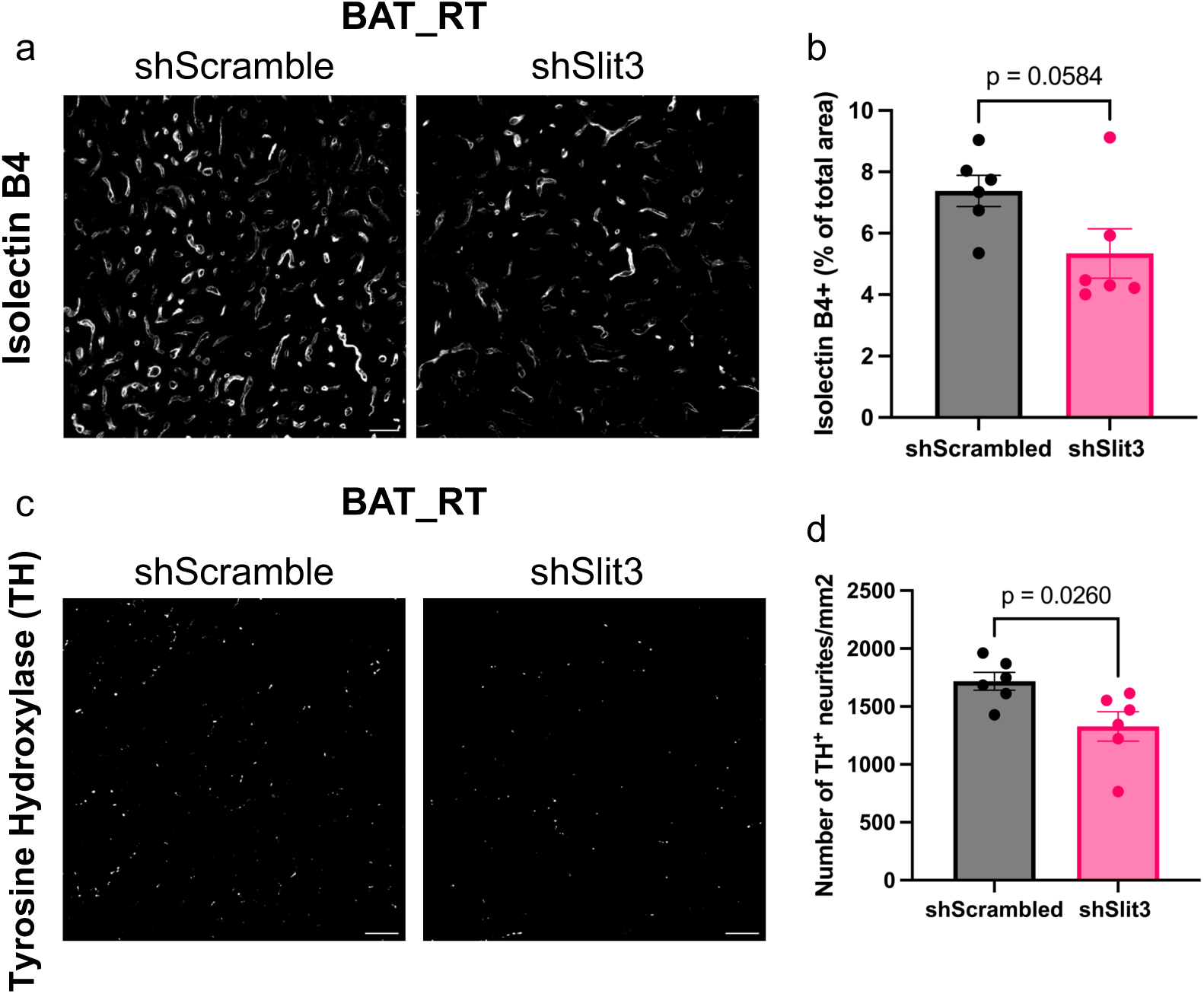
(a) Representative images of Isolectin B4 staining and (b) quantification of the percentage of Isolectin B4+ area in BAT of mice receiving AAV-shSlit3 or scramble shRNA and housed at room temperature. Scale bar = 20 µm. (c) Representative images of TH staining and (d) quantification of the number of TH+ neurites per area in BAT of mice receiving AAV-shSlit3 or scramble shRNA and housed at room temperature. Scale bar = 25 µm. N = 6 per group. Data are presented as means ± SEM and analyzed by unpaired two-sided Student’s t-tests (b and d)

**Supplementary Figure 8. Related to Figure 2.**
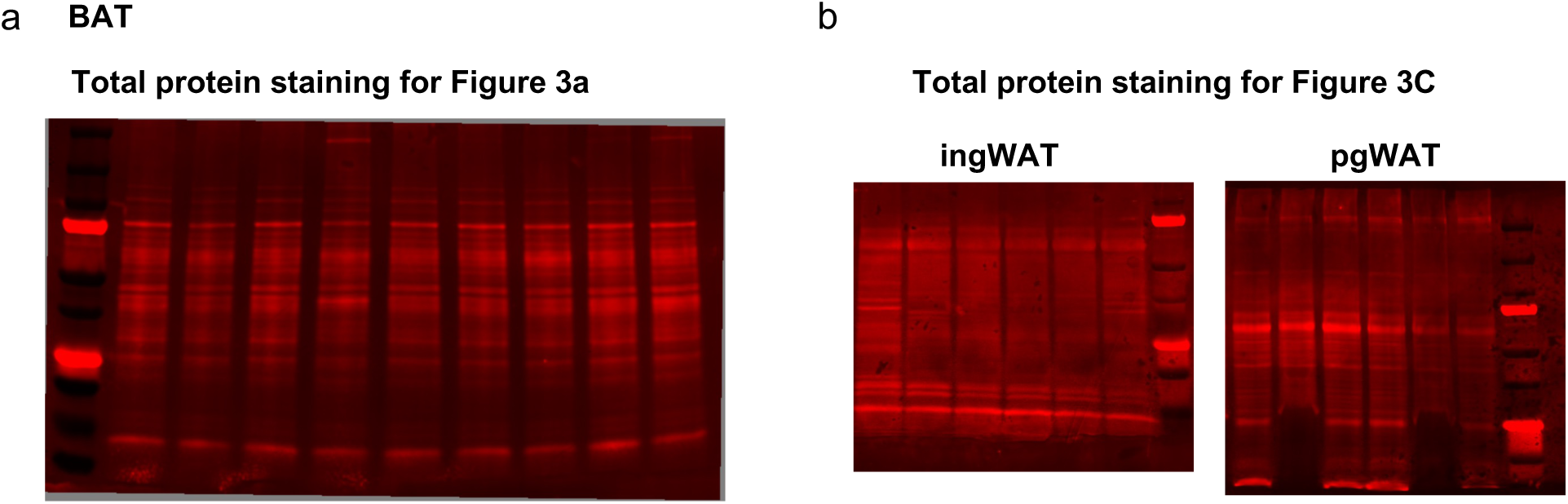
(a-b) Total protein staining corresponding to Figures 3a and 3c.

**Supplementary Figure 9. Related to Figure 3.**
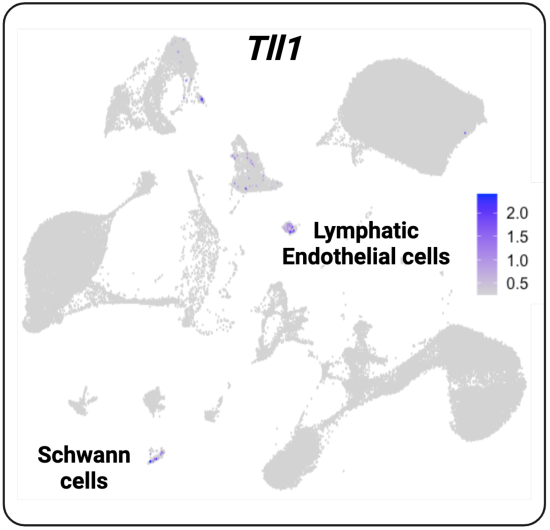
Expression of *Tll1* in scRNA-seq data of mouse BAT.

**Supplementary Figure 10. Related to Figure 5.**
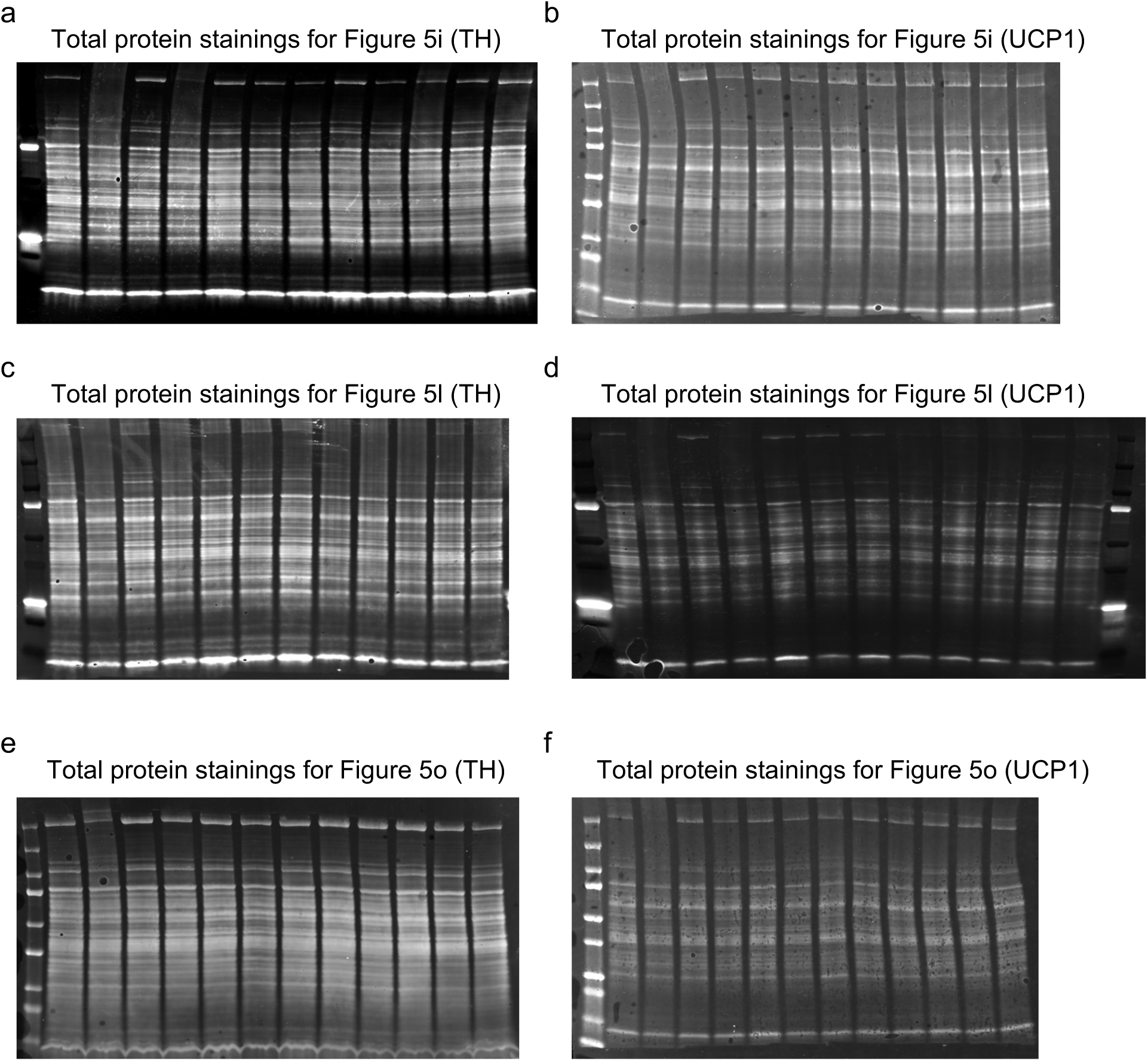
(a–f) Total protein staining corresponding to Figures 5i, 5l, and 5o.

**Supplementary Figure 11. Related to Figure 7.**
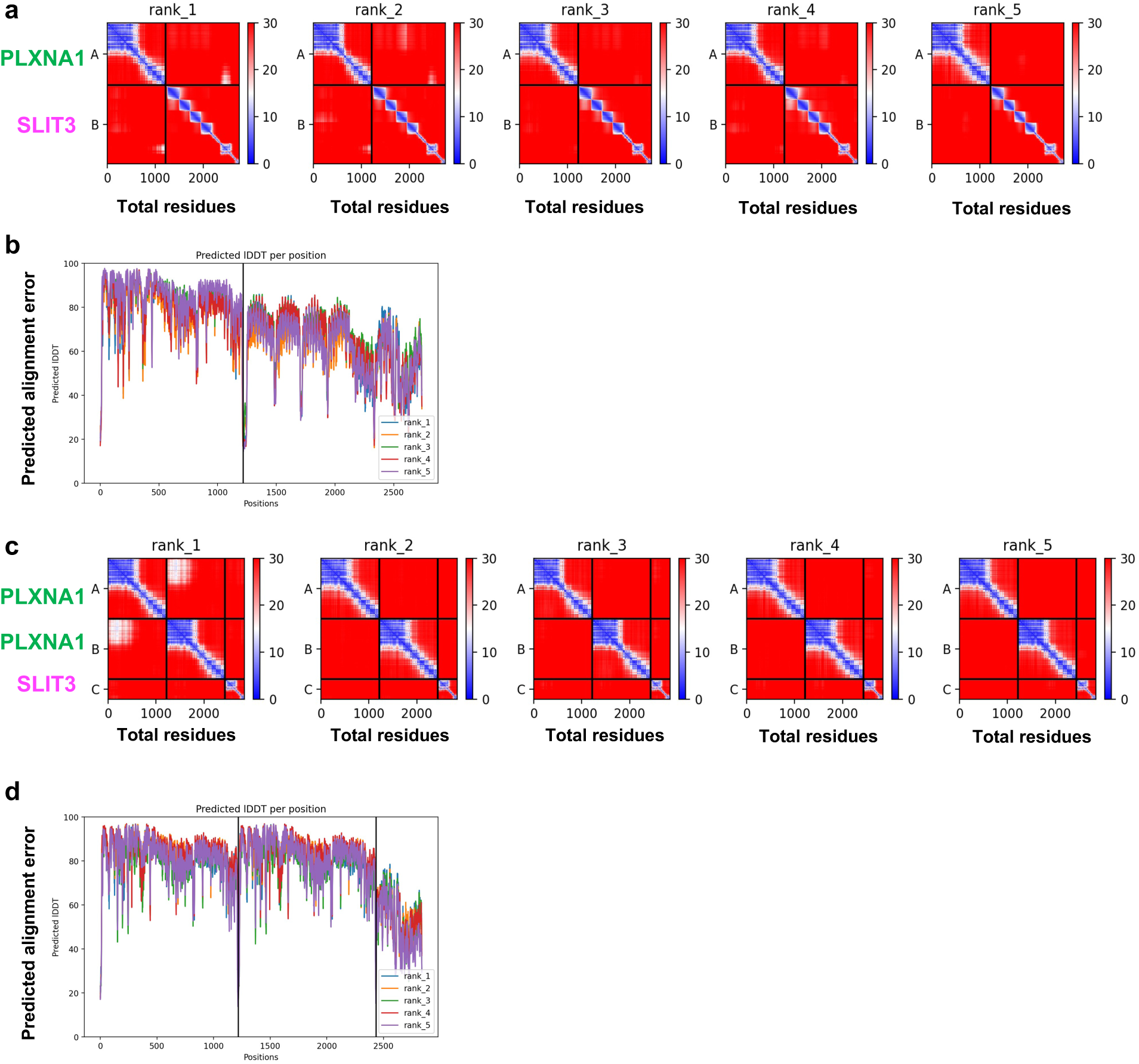
AlphaFold2 Multimer predicts the human PLXNA1-SLIT3 complex. Diagnostic plots and images of 3D models demonstrate the quality of the protein complex predictions using the sequences of human PLXNA1 extracellular domains 1-10 and the C-terminal region of the human SLIT3. (a) The PAE (predicted error in A° between every pair of residues) plots of monomeric PLXNA1 and SLIT3 interaction model. A and B tiles represent PLXNA1and Slit3, respectively. The X, Y axes indicate the total amino acids in these analyses. (b) The pLDDT (predicted Local Distance Difference between all atoms) plots show the predicted IDDT per residue in the top five ranked models for monomeric PLXNA1- SLIT3 complex. The X, Y axes indicate the total amino acids in these analyses. (c) The PAE plot of dimeric PLXNA1 and monomeric SLIT3 interaction model. A and B tiles represent each PLXNA1 monomer, and the C tile shows SLIT3. (d) The pLDDT plots show the predicted IDDT per residue in the top five ranked models for the PLXNA1 dimer complex with SLIT3.

**Supplementary Figure 12. Related to Figure 7.**
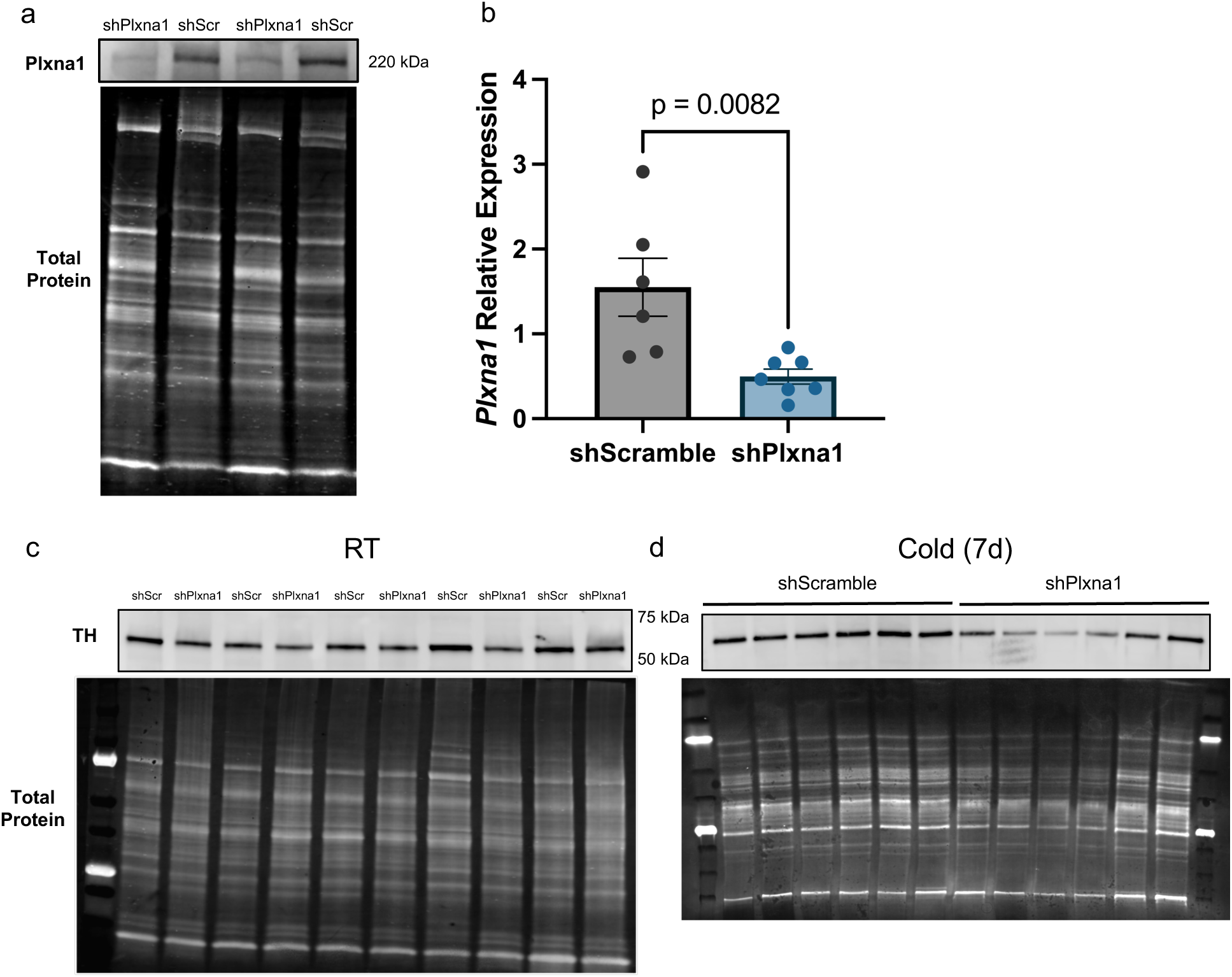
(a) PLXNA1 protein and (b) transcript expression levels. (c) Total TH protein levels in BAT of mice receiving AAV-shPlxna1 or scramble shRNA. N = 5 per group. Data are presented as mean ± SEM and analyzed by unpaired two-sided Student’s t-test (b).

## References

1 Cannon, B. & Nedergaard, J. Brown adipose tissue: function and physiological significance. Physiol Rev 84, 277–359, doi:10.1152/physrev.00015.2003 (2004).

2 Shamsi, F., Wang, C. H. & Tseng, Y. H. The evolving view of thermogenic adipocytes - ontogeny, niche and function. Nat Rev Endocrinol 17, 726–744, doi:10.1038/s41574-021-00562-6 (2021).

3 Xue, Y. et al. Hypoxia-independent angiogenesis in adipose tissues during cold acclimation. Cell Metab 9, 99–109, doi:10.1016/j.cmet.2008.11.009 (2009).

4 Takahashi, A., Shimazu, T. & Maruyama, Y. Importance of sympathetic nerves for the stimulatory effect of cold exposure on glucose utilization in brown adipose tissue. Jpn J Physiol 42, 653–664, doi:10.2170/jjphysiol.42.653 (1992).

5 Shamsi, F. et al. Vascular smooth muscle-derived Trpv1(+) progenitors are a source of cold-induced thermogenic adipocytes. Nat Metab 3, 485–495, doi:10.1038/s42255-021-00373-z (2021).

6 Shamsi, F., Zheng, R., Ho, L. L., Chen, K. & Tseng, Y. H. Comprehensive analysis of intercellular communication in the thermogenic adipose niche. Commun Biol 6, 761, doi:10.1038/s42003-023-05140-2 (2023).

7 Cannavino, J. & Gupta, R. K. Mesenchymal stromal cells as conductors of adipose tissue remodeling. Genes Dev 37, 781–800, doi:10.1101/gad.351069.123 (2023).

8 Wang, Y. N. et al. Slit3 secreted from M2-like macrophages increases sympathetic activity and thermogenesis in adipose tissue. Nat Metab 3, 1536–1551, doi:10.1038/s42255-021-00482-9 (2021).

9 Emont, M. P. et al. A single-cell atlas of human and mouse white adipose tissue. Nature 603, 926–933, doi:10.1038/s41586-022-04518-2 (2022).

10 Lin, Z., Chi, J. & Cohen, P. A Clearing Method for Three-Dimensional Imaging of Adipose Tissue. Methods Mol Biol 2448, 73–82, doi:10.1007/978-1-0716-2087-8_4 (2022).

11 Xu, R. et al. Targeting skeletal endothelium to ameliorate bone loss. Nat Med 24, 823–833, doi:10.1038/s41591-018-0020-z (2018).

12 Kang, S. H., Fukaya, M., Yang, J. K., Rothstein, J. D. & Bergles, D. E. NG2+ CNS glial progenitors remain committed to the oligodendrocyte lineage in postnatal life and following neurodegeneration. Neuron 68, 668–681, doi:10.1016/j.neuron.2010.09.009 (2010).

13 Brose, K. et al. Slit proteins bind Robo receptors and have an evolutionarily conserved role in repulsive axon guidance. Cell 96, 795–806, doi:10.1016/s0092-8674(00)80590-5 (1999).

14 Choi, C. H. J. et al. LRG1 is an adipokine that promotes insulin sensitivity and suppresses inflammation. Elife 11, doi:10.7554/eLife.81559 (2022).

15 Kellermeyer, R. et al. Proteolytic cleavage of Slit by the Tolkin protease converts an axon repulsion cue to an axon growth cue in vivo. Development 147, doi:10.1242/dev.196055 (2020).

16 Talantikite, M. et al. Inhibitors of BMP-1/tolloid-like proteinases: efficacy, selectivity and cellular toxicity. FEBS Open Bio 8, 2011–2021, doi:10.1002/2211-5463.12540 (2018).

17 Delloye-Bourgeois, C. et al. PlexinA1 is a new Slit receptor and mediates axon guidance function of Slit C-terminal fragments. Nat Neurosci 18, 36–45, doi:10.1038/nn.3893 (2015).

18 Nguyen Ba-Charvet, K. T., et al. Diversity and specificity of actions of Slit2 proteolytic fragments in axon guidance. J Neurosci 21, 4281–4289 (2001).

19 Watanabe, T., Watanabe-Kominato, K., Takahashi, Y., Kojima, M. & Watanabe, R. Adipose Tissue-Derived Omentin-1 Function and Regulation. Compr Physiol 7, 765–781, doi:10.1002/cphy.c160043 (2017).

20 Yang, R. Z. et al. Identification of omentin as a novel depot-specific adipokine in human adipose tissue: possible role in modulating insulin action. Am J Physiol Endocrinol Metab 290, E1253–1261, doi:10.1152/ajpendo.00572.2004 (2006).

21 Amano, S. et al. Bone morphogenetic protein 1 is an extracellular processing enzyme of the laminin 5 gamma 2 chain. J Biol Chem 275, 22728–22735, doi:10.1074/jbc.M002345200 (2000).

22 Kessler, E., Takahara, K., Biniaminov, L., Brusel, M. & Greenspan, D. S. Bone morphogenetic protein-1: the type I procollagen C-proteinase. Science 271, 360–362, doi:10.1126/science.271.5247.360 (1996).

23 Hopkins, D. R., Keles, S. & Greenspan, D. S. The bone morphogenetic protein 1/Tolloid- like metalloproteinases. Matrix Biol 26, 508–523, doi:10.1016/j.matbio.2007.05.004 (2007).

24 Muir, A. M. et al. Induced ablation of Bmp1 and Tll1 produces osteogenesis imperfecta in mice. Hum Mol Genet 23, 3085–3101, doi:10.1093/hmg/ddu013 (2014).

25 Zhou, Y., Gunput, R.-A. F. & Pasterkamp, R. J. Semaphorin signaling: progress made and promises ahead. Trends in Biochemical Sciences 33, 161–170, 10.1016/j.tibs.2008.01.006 (2008).

26 Alto, L. T. & Terman, J. R. Semaphorins and their Signaling Mechanisms. Methods Mol Biol 1493, 1–25, doi:10.1007/978-1-4939-6448-2_1 (2017).

27 Guilherme, A., Henriques, F., Bedard, A. H. & Czech, M. P. Molecular pathways linking adipose innervation to insulin action in obesity and diabetes mellitus. Nat Rev Endocrinol 15, 207–225, doi:10.1038/s41574-019-0165-y (2019).

28 Jiang, H., Ding, X., Cao, Y., Wang, H. & Zeng, W. Dense Intra-adipose Sympathetic Arborizations Are Essential for Cold-Induced Beiging of Mouse White Adipose Tissue. Cell Metab 26, 686–692 e683, doi:10.1016/j.cmet.2017.08.016 (2017).

29 Murano, I., Barbatelli, G., Giordano, A. & Cinti, S. Noradrenergic parenchymal nerve fiber branching after cold acclimatisation correlates with brown adipocyte density in mouse adipose organ. J Anat 214, 171–178, doi:10.1111/j.1469-7580.2008.01001.x (2009).

30 Bougneres, P. et al. In vivo resistance of lipolysis to epinephrine. A new feature of childhood onset obesity. J Clin Invest 99, 2568–2573, doi:10.1172/JCI119444 (1997).

31 Heinonen, S. et al. Impaired Mitochondrial Biogenesis in Adipose Tissue in Acquired Obesity. Diabetes 64, 3135–3145, doi:10.2337/db14-1937 (2015).

32 Guo, T. et al. Adipocyte ALK7 links nutrient overload to catecholamine resistance in obesity. Elife 3, e03245, doi:10.7554/eLife.03245 (2014).

33 Pirzgalska, R. M. et al. Sympathetic neuron-associated macrophages contribute to obesity by importing and metabolizing norepinephrine. Nat Med 23, 1309–1318, doi:10.1038/nm.4422 (2017).

34 Saxton, S. N., Withers, S. B. & Heagerty, A. M. Emerging Roles of Sympathetic Nerves and Inflammation in Perivascular Adipose Tissue. Cardiovasc Drugs Ther 33, 245–259, doi:10.1007/s10557-019-06862-4 (2019).

35 Daily, J. W., Liu, M. & Park, S. High genetic risk scores of SLIT3, PLEKHA5 and PPP2R2C variants increased insulin resistance and interacted with coffee and caffeine consumption in middle-aged adults. Nutr Metab Cardiovasc Dis 29, 79–89, doi:10.1016/j.numecd.2018.09.009 (2019).

36 Liu, Y. J. et al. Genome-wide association scans identified CTNNBL1 as a novel gene for obesity. Hum Mol Genet 17, 1803–1813, doi:10.1093/hmg/ddn072 (2008).

37 Baca, P. et al. DNA methylation and gene expression analysis in adipose tissue to identify new loci associated with T2D development in obesity. Nutr Diabetes 12, 50, doi:10.1038/s41387-022-00228-w (2022).

38 van der Klaauw, A. A. et al. Human Semaphorin 3 Variants Link Melanocortin Circuit Development and Energy Balance. Cell 176, 729–742 e718, doi:10.1016/j.cell.2018.12.009 (2019).

39 Shamsi, F. et al. FGF6 and FGF9 regulate UCP1 expression independent of brown adipogenesis. Nat Commun 11, 1421, doi:10.1038/s41467-020-15055-9 (2020).

40 Berry, R. et al. Imaging of adipose tissue. Methods Enzymol 537, 47–73, doi:10.1016/b978-0-12-411619-1.00004-5 (2014).

41 Mina, A. I. et al. CalR: A Web-Based Analysis Tool for Indirect Calorimetry Experiments. Cell Metab 28, 656–666.e651, doi:10.1016/j.cmet.2018.06.019 (2018).

42 Chi, J., Crane, A., Wu, Z. & Cohen, P. Adipo-Clear: A Tissue Clearing Method for Three- Dimensional Imaging of Adipose Tissue. J Vis Exp, doi:10.3791/58271 (2018).

43 Jumper, J. et al. Highly accurate protein structure prediction with AlphaFold. Nature 596, 583–589, doi:10.1038/s41586-021-03819-2 (2021).

44 Evans, R., et al. Protein complex prediction with AlphaFold-Multimer. *bioRxiv*, 2021.2010.2004.463034, doi:10.1101/2021.10.04.463034 (2022).

45 Steinegger, M. & Soding, J. MMseqs2 enables sensitive protein sequence searching for the analysis of massive data sets. Nat Biotechnol 35, 1026–1028, doi:10.1038/nbt.3988 (2017).

46 Mirdita, M. et al. ColabFold: making protein folding accessible to all. Nat Methods 19, 679–682, doi:10.1038/s41592-022-01488-1 (2022).

47 Pettersen, E. F. et al. UCSF Chimera--a visualization system for exploratory research and analysis. J Comput Chem 25, 1605–1612, doi:10.1002/jcc.20084 (2004).

48 Langhardt, J. et al. Effects of Weight Loss on Glutathione Peroxidase 3 Serum Concentrations and Adipose Tissue Expression in Human Obesity. Obes Facts 11, 475–490, doi:10.1159/000494295 (2018).

49 Mardinoglu, A. et al. Extensive weight loss reveals distinct gene expression changes in human subcutaneous and visceral adipose tissue. Scientific Reports 5, 14841, doi:10.1038/srep14841 (2015).

50 Blüher, M. Metabolically Healthy Obesity. Endocr Rev 41, doi:10.1210/endrev/bnaa004 (2020).

51 Klöting, N. et al. Insulin-sensitive obesity. Am J Physiol Endocrinol Metab 299, E506–515, doi:10.1152/ajpendo.00586.2009 (2010).

52 Picelli, S. et al. Full-length RNA-seq from single cells using Smart-seq2. Nat Protoc 9, 171–181, doi:10.1038/nprot.2014.006 (2014).

53 Hagemann, T. et al. Laminin α4 Expression in Human Adipose Tissue Depots and Its Association with Obesity and Obesity Related Traits. Biomedicines 11, doi:10.3390/biomedicines11102806 (2023).

54 Bray, N. L., Pimentel, H., Melsted, P. & Pachter, L. Near-optimal probabilistic RNA-seq quantification. Nat Biotechnol 34, 525–527, doi:10.1038/nbt.3519 (2016).

